# Processive activity of the replicative DNA polymerases in the replisome of live eukaryotic cells

**DOI:** 10.1101/780270

**Authors:** Nitin Kapadia, Ziad W. El-Hajj, Huan Zheng, Thomas R. Beattie, Angela Yu, Rodrigo Reyes-Lamothe

**Affiliations:** Department of Biology, McGill University, 3649 Sir William Osler, Montreal, Qc, H3G 0B1 Canada

## Abstract

DNA replication is carried out by a multi-protein machine called the replisome. In *Saccharomyces cerevisiae*, the replisome is composed of over 30 different proteins arranged into multiple subassemblies, each performing distinct activities. Synchrony of these activities is required for efficient replication and preservation of genomic integrity. How this is achieved is particularly puzzling at the lagging strand, where current models of the replisome architecture propose turnover of the canonical lagging strand polymerase, Pol δ, at every cycle of Okazaki fragment synthesis. Here we established single-molecule fluorescence microscopy protocols to study the binding kinetics of individual replisome subunits in live *S. cerevisiae*. Our results show long residence times for most subunits at the active replisome, supporting a model where all subassemblies bind tightly and work in a coordinated manner for extended periods, including Pol δ, hence redefining the architecture of the active eukaryotic replisome.

## INTRODUCTION

A crucial step during cell proliferation is the duplication of the genome, composed of long DNA molecules. DNA replication is carried out by a specialized multiprotein machine called the replisome. In all organisms, the replisome accomplishes a few basic functions: progressive unwinding of the double-stranded DNA, the synthesis of an RNA molecule that serves as primer, and the polymerization of DNA. In eukaryotic cells, DNA unwinding is catalysed by the CDC45/MCM2-7/GINS (CMG) helicase. Three replicative DNA polymerases act at the replisome: Pol α, Pol ε and Pol δ. Pol α carries out the RNA primer synthesis at the lagging strand – proceeding in an opposite direction to DNA unwinding – and extends primers using its DNA polymerase activity. Pol ε and Pol δ synthesize most of the DNA on the leading and lagging strands, respectively. Whilst Pol ε is thought to interact stably with CMG, Pol δ does not appear to have a stable interaction with the rest of the replisome (Schauer and O’Donnell, 2017; Sengupta et al., 2013). Hence, each active copy of Pol δ is assumed to have a limited processivity, synthesizing the length of an Okazaki fragment (∼160bp), before unbinding from DNA (Bell and Labib, 2016; Smith and Whitehouse, 2012). In addition to these basic functions, the replisome contains multiple other proteins serving structural or other specialized roles, such as RPA, which binds to single-stranded DNA (ssDNA), and Ctf4, which is thought to act as a recruiting hub for Pol α and other proteins (Figure 3A) (reviewed in (Bell and Labib, 2016)).

It is thought that a stable architecture of the replisome helps ensure that the basic activities occur in an orderly and synchronized manner, thus limiting the accumulation of ssDNA near the sites of replication, which can lead to increased mutation rates and to breaks in DNA (Lopes et al., 2006; Yang et al., 2008). However, plasticity of the architecture over time – through dissociation of subunits – has been observed in bacteria (Beattie et al., 2017; Lewis et al., 2017; Liao et al., 2016). This leads to the question of how this balance of synchronized work and dynamic behaviour of individual subunits is achieved in the eukaryotic replisome. *In vitro* studies of the binding kinetics of individual subunits of the replisome have been limited in the past by the great number of components and the presence of post-translational modifications, both of which can influence the strength of interaction between proteins and replisome function.

Here we established a single-molecule fluorescence microscopy approach to study binding kinetics of nuclear proteins in live budding yeast and used Fluorescence Recovery After Photobleaching (FRAP) to validate it. We used this approach to characterize the binding kinetics of eukaryotic replisome subunits in live cells. Our results show that a single copy of Pol α remain at the replication fork for multiple rounds of Okazaki fragment synthesis. And that individual copies of Pol ε, and surprisingly, of Pol δ remain at the replication fork for an average time of at least 5 minutes, suggesting a different architecture for the eukaryotic replisome than the one generally accepted.

## RESULTS

### Single-molecule tracking of nuclear proteins in live yeast

We first established a set of experimental and analysis protocols for the use of single-molecule fluorescence microscopy in budding yeast. In order to specifically label a protein of interest (POI), we genetically tagged the POI with HaloTag, obtaining haploid strains carrying fusions to single-copies of the genes at their native chromosomal locus(Los et al., 2008)(Figure S1-S3; Supplemental Text). Use of HaloTag allowed us to utilize organic fluorophores brighter than currently available fluorescent proteins, facilitating single-molecule detection. Halo-tagged POIs were visualized by incubating cells with a low concentration of PAJF549, a cell-permeable bright photoactivatable dye, which was conjugated to the HaloTag substrate (Figure 1A) (Grimm et al., 2016). The result of this treatment is the covalent binding of PAJF549 to the Halo-POI. Labelled cells were transferred to an agarose pad containing SC medium on a microscope slide.

**Figure 1.**
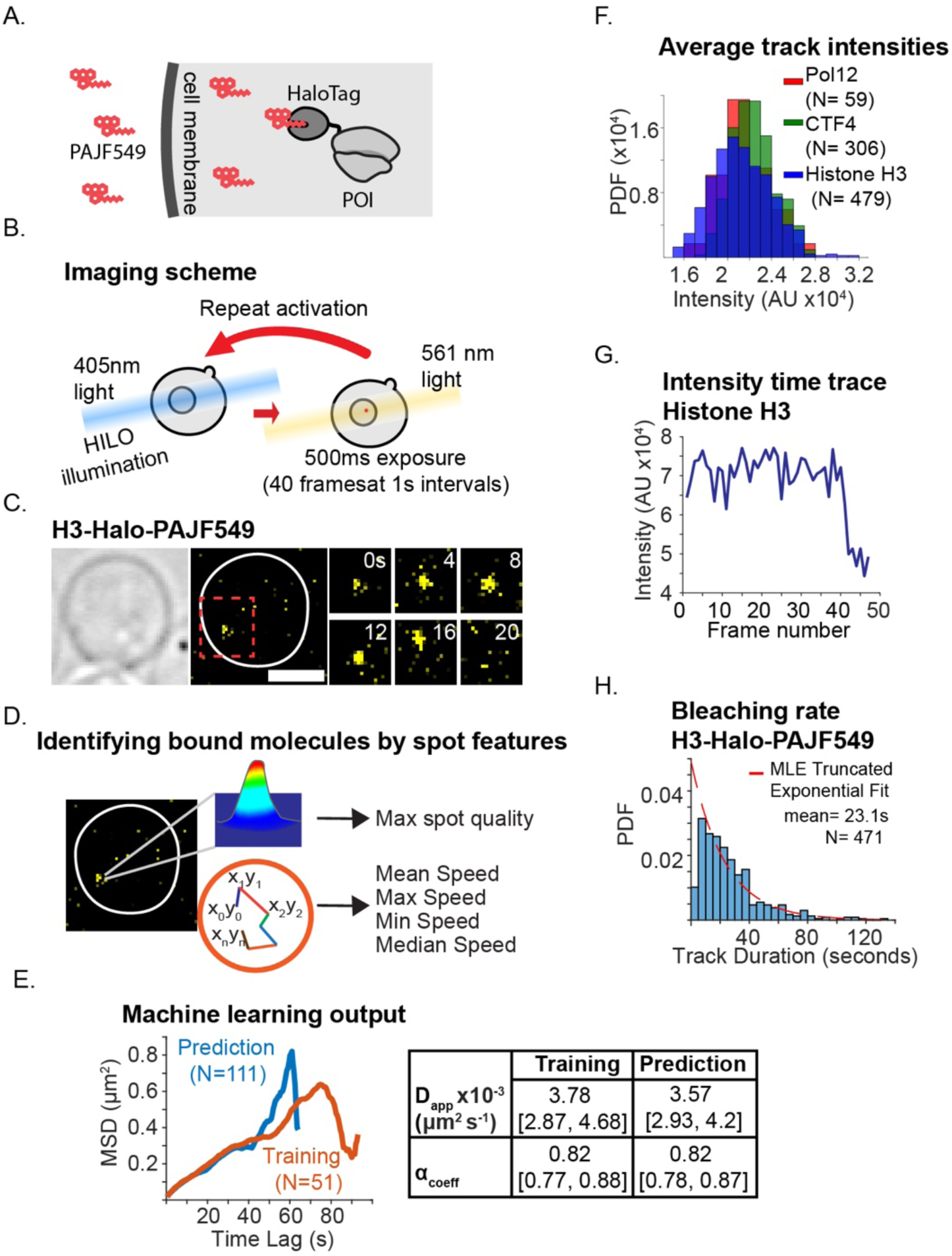
Single-molecule tracking in the yeast nucleus. (**A**) Strains carrying a HaloTag fusion to our protein of interest (POI). We incubated them with a PAJF549 dye + SC media solution to allow some of the dye to be incorporated into the cell, and subsequently into the nucleus. (**B**) We exposed Halo-tagged yeast cells to a single exposure of low-intensity 405nm light, resulting in the stochastic activation of few molecules per nucleus. This was followed by time-lapse imaging using Highly Inclined and Laminated Optical sheet (HILO) illumination and 561nm light for fluorescence detection. Multiple rounds of activation and imaging were performed to increase sample size. (**C**) Representative images of single-molecules of H3-Halo-PAJF549 in live cells. The image on the left shows the entire cell. Images on the right show a magnified view of the square drawn on the left image, and show individual timepoints over 20 seconds. Scale bar 2 μm. (**D**) Information on the intensity and speed of single molecules of H3-Halo-PAJF549 was extracted and used to identify chromatin-bound molecules of replisome subunits. (**E**) MSD analysis of the classification outcome from the machine-learning algorithm used to identify chromatin-bound molecules. We isolated tracks manually classified as being bound from the training data set and calculated an averaged MSD curve, to determine their diffusive behaviour (see Materials and Methods). We then performed the same calculation on tracks classified as being bound with our ML approach, from a different data set that was collected under same acquisition settings as the original training data. (**F**) Distributions of average track intensities after GMM fitting and clustering to isolate tracks representing single-molecules. (**G**) Representative intensity-time trace from H3-Halo, showing a noticeable single-step with a similar intensity value as the intensity of spots selected by the ML approach. (**H**) Characteristic bleaching rate for the fluorophore PAJF549 using H3-Halo as control, whose molecules should bleach before unbinding.

The imaging protocol used to track single-molecules was similar to that previously described for *Escherichia coli* (Beattie et al., 2017), where a single exposure to a low dose of 405nm light, to activate few molecules of PAJF549 per cell, was followed by multiple events of 561nm excitation-light at fixed intervals to track the activated molecules. However, here we did multiple rounds of activation-imaging to maximize the sample size (Figure 1B). Camera integration times of 500 milliseconds resulted in the motion-blurring of diffusive molecules, facilitating the tracking of chromatin-bound molecules represented as foci. All our imaging was done using HILO illumination to increase the signal-to-noise ratio (Tokunaga et al., 2008). In addition, to minimize light refraction caused by the cell wall (Smith et al., 2015), we did our experiments in the presence of 30% of the refractive media-matching Optiprep (Figure S4)(Boothe et al., 2017).

We used a strain carrying a fusion of the histone H3 to HaloTag (H3-Halo) (Ball et al., 2016; Los et al., 2008) as a control, due to the expected long-residence binding of H3 on chromatin – in mammalian cells this estimate is of several hours (Kimura and Cook, 2001). To increase the confidence in our results, we only included tracks where molecules were localized for at least four frames; we used a truncated exponential model to correct for tracks shorter than this minimal length. Analysis of experiments of a strain lacking a HaloTag fusion, but incubated with the dye, resulted in the detection of no spots, showing that non-specific dye binding does not represent a source of noise in our experiments (Figure S5). Using H3-Halo we characterized the rate of bleaching of the fluorophore, the localization linking distance for tracking chromatin-bound proteins in our timelapses, and the traits of a single-fluorescent molecule in our system, which we exploited for robust classification using a machine-learning (ML) approach (Figure 1C-E,H). Unbiased identification of chromatin-bound single-molecules was done using a random forest-based machine learning algorithm, which used multiple traits of molecular movement and intensity for classification (Figure 1D-E; Figure S6).

Our experimental and analysis protocols were designed to exclusively incorporate in our results, data from single-molecules: we used low dye concentrations to label only a fraction of proteins; we used low laser activation power to activate few dye molecules per cell; we discarded tracks found in regions of high density (Figure S7); and we used a Gaussian mixture model (GMM) fitting to the mean track intensities to discard bright spots likely representing multiple molecules (Figure S8). We expected that spots of single fluorescent molecules should bleach in a single step, and that their intensities should distribute normally. Indeed, single-bleaching steps can be observed in the data from histone H3-Halo (Figure 1F; Figure S9). The values of these single steps correlated with the peak for the mean track intensities for H3-Halo, and remarkably, it also correlated with the mean track intensities of replisome subunits (Figure 1G). Taken together, these results provide support that our approach selectively analysed tracks representing single-molecules.

### Detection of single subunits in active replisomes

We then sought to confirm that spots observed in strains carrying Halo-tagged replisome subunits represented proteins engaged in active replisomes. To test if spots from Halo-tagged replisome subunits localize at replication forks we used a PCNA-mNeonGreen fusion (PCNA-mNG)(Shaner et al., 2013). PCNA has been reported to be a reliable marker of S phase by the presence of distinct foci (Figure S10) (Kitamura et al., 2006), which in addition we exploited during image analysis by segmenting nuclei containing heterogeneous intensity (Figure 2F). We tracked cells that carried both PCNA-mNG and a Halo-tagged DNA polymerase subunit at intervals of 1 or 2 minutes for at least 30 minutes (Figure 2A-B; Figure S11). In these experiments photoactivation was higher than in single-molecule experiments to increase the number of fluorescent molecules and pictures were taken using a single focal plane. In addition, spots in consecutive frames were not linked into tracks. We observed 22-35% of nuclei in S phase cells carried HaloTag spots, against 4-7% in non-S phase cells (Figure 2C). Spots representing DNA polymerases during S phase often co-localized with PCNA spots (Figure 2A-B; Figure S11) and their frequency increased with the number of replisomes, peaking at mid S phase (Figure 2D-E).

**Figure 2.**
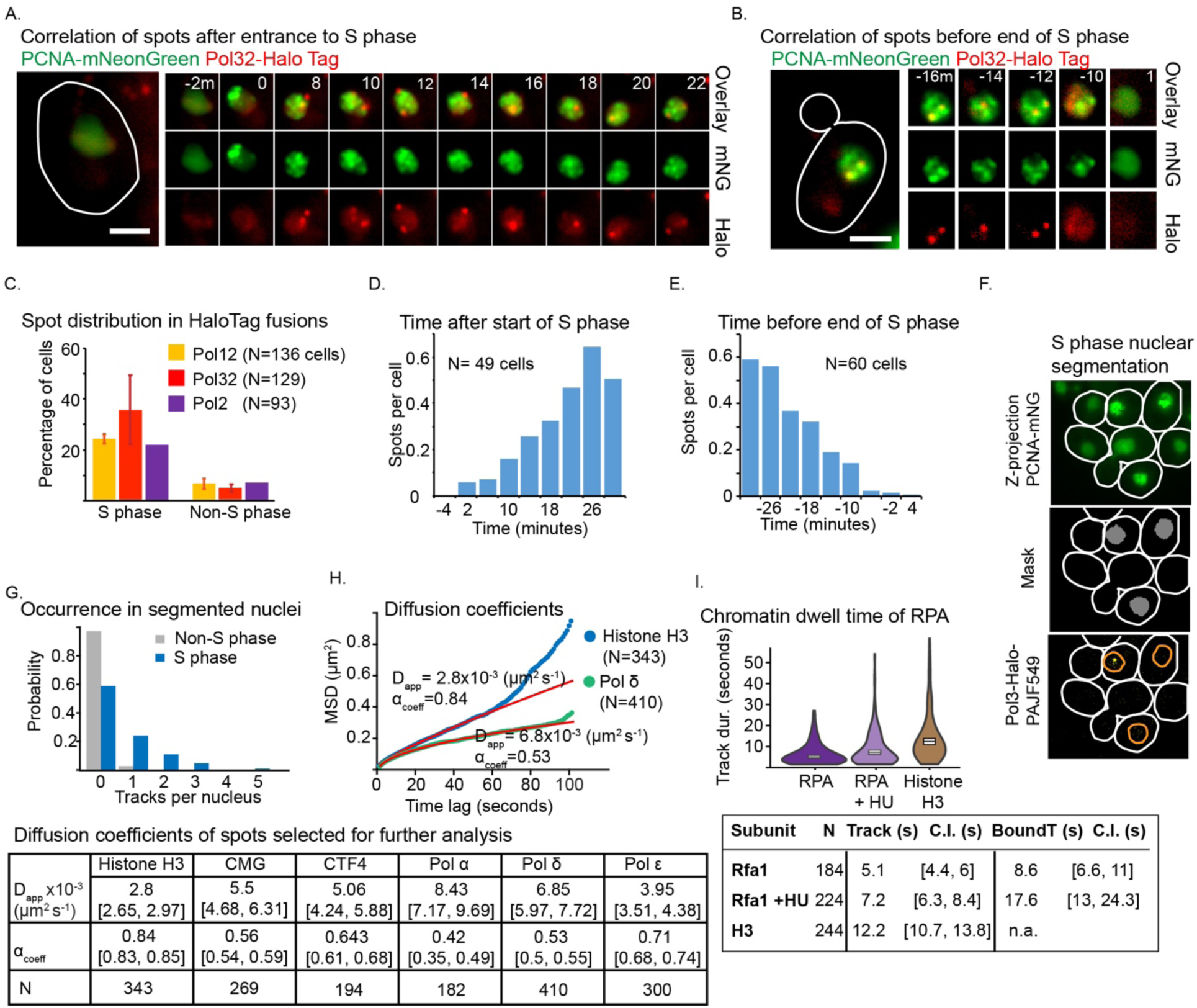
Study of replisome subunits in active replisomes. (**A**) Representative images of time lapses of Pol32-Halo and PCNA-mNG showing a cell entering S phase. Time points not shown had no Pol32-Halo foci. 0 time point shows the appearance of PCNA-mNG spots. Scale bar 2μm. (**B**) Representative images of the same strain as in A showing a cell exiting S phase. Time points not shown had no Pol32-Halo foci. 0 time point shows the disappearance of PCNA-mNG spots. Scale bar 2μm. (**C**) Frequency of detecting overactivated Halo-tagged polymerase spots in S vs non-S phase cells. Nuclei were segmented based on the heteogeneity of PCNA-mNeonGreen intensities (see panel F). Halo-tagged molecules were overactivated to allow detection of multiple spots per cell, and the spots were classified based in their appearance in S or non-S nuclei. (**D**) Increase in number of detected Pol-Halo spots as cells enter and proceed towards the peak of S phase. (**E**) Decrease in number of detected Pol-Halo spots as cells move from mid-S to the end of S phase. (**F**) Representative image of the procedure followed to segment nuclei of cells undergoing S phase. Smooth Manifold Extraction (SME) was used to project the intensity of a Z-stack of PCNA-mNeonGreen covering the whole cell while preserving spot-like features. S phase nuclei were selected based on high heterogeneity in the distribution of intensities. (**G**) Representative diagram showing the tracks found in the nucleus after ML classification in non S -phase (N =187 nuclei) vs S phase (N=162), from Pol32 data. (**H**) Top) Comparison of diffusive behaviour between Histone H3 and Pol δ. Note that the tracks in Histone H3 were taken from cells at varying stages of the cell cycle while those of Pol δ were in S phase cells only. Bottom) Table showing diffusive properties of replisomal proteins. (**I**) Results of RPA and RPA with 0.2M HU. Data was collected with 500ms exposure and interval time between consecutive pictures.

We performed a similar analysis for our single-molecule experiments using Pol32-Halo and 1-second intervals between pictures. Here, after ML classification, we detected tracks in >40% of cells categorized as S phase, compared with <5% in non-S phase cells (Figure 2G). The small number of tracks in cells not categorized as S phase in our single-molecule experiments may be explained by multiple reasons: errors in our S phase categorization scheme – for which we used stringent parameters that are prone to exclude some S phase cells – errors in the ML classification approach; and Halo-tagged proteins bound to chromatin but not at replication forks. Although we acknowledge the latter possibility, the tracks in these cells had a mean duration of ∼6s, which as will be apparent in later sections, is much shorter than the durations of those found in S phase cells. Therefore, the great majority of the tracks used should represent proteins bound at active replisomes.

If the molecules that produce these fluorescent spots are indeed bound to replisomes they should diffuse similarly to other proteins bound to chromatin. To test this, we estimated the diffusive properties of Halo-tagged proteins in our single-molecule experiments using 1-second intervals. We found not only that the values were consistent among the different proteins – ranging from 2-8×10-3 μm s-1 – but also that the diffusion coefficients and anomalous diffusion constants were similar to those reported for chromosomal loci, indicating that the spots detected represent chromatin-bound molecules (Figure 2H; Figure S12) (Hajjoul et al., 2013). This illustrates that our experimental design, in combination with our ML approach, is robust enough to detect predominantly chromatin-bound proteins, supporting the notion that spots for Halo-tagged replisome subunits represent copies forming part of replisomes.

How can we be more confident that the molecules we detect represent proteins engaged in active replisomes? To answer this question, we tested whether our single-molecule experiments could report a change in the dynamics of the replisome after a perturbation of its activity. We used a subunit Halo-tagged version of the single-stranded binding protein RPA, Rfa1. During active DNA replication, considering a fork progression rate of 1.6kb/min and an Okazaki fragment size of 160bp (Sekedat et al., 2010; Smith and Whitehouse, 2012), we predicted that Rfa1 should have a residence time of ∼6s at the replication fork. We tracked Rfa1-Halo spot lifetimes using 0.5-second intervals between pictures. The distribution of track durations was fitted to a truncated exponential model – as done above for H3-Halo – from which we obtained a time constant of 5.1 seconds. After correcting for bleaching, we estimated a residence time of RPA to be 8.6s, consistent with the predicted residence time (Figure 2I). We expected that treatment with hydroxyurea (HU) would cause a slowdown of replisomes due to dNTP depletion (Alvino et al., 2007), resulting in a longer residence time for RPA. Indeed, we estimated a track duration of 7.2 seconds and a residence time of 17.6 seconds for Rfa1-Halo when treated with 0.2M HU. The remaining slower exchange rate observed probably reflects residual DNA replication and concentration-dependent exchange of RPA (Gibb et al., 2014). Overall, the results above indicate that our system can be used to study the dynamics of proteins engaged in active replisomes.

### Replisome subunits are stably bound

We applied these newly developed strategies to evaluate the residence times of multiple replisome subunits. We expected long residence times for the CMG helicase and the leading-strand Pol ε. In contrast, we expected much shorter binding for Pol δ, consistent with the use of a different copy of the polymerase for the synthesis of each Okazaki fragment. We performed multiple experiments for each subunit to check for consistency, but given the small sample size per experiment due to the sparse activation of PAJF549, leading to greater variability in the individual estimates, we combined the track durations across multiple experiments and then performed a fit, to obtain a final estimate. Using 1-second intervals in a single-imaging plane, we observed lifetimes of fluorescent foci that were indistinguishable from H3-Halo (our bleaching control) in cells carrying Cdc45-Halo and Mcm4-Halo (Figure 3B-C; Table S6, S8), consistent with a reported tight binding of CMG at chromatin (Yeeles et al., 2015). Similarly, Pol ε (Pol2-Halo and Dpb4-Halo) and the Ctf4 subunit – both of which have a direct interaction with the CMG helicase (Sengupta et al., 2013; Simon et al., 2014) – exhibited lifetimes similar to those of the CMG complex, suggesting they interact tightly with it. Unexpectedly, Pol3-Halo and Pol32-Halo, two subunits of Pol δ, also showed long track durations indicative of stable binding (Figure 3C; Video S1). This was surprising since there is no reported connection between Pol δ and the CMG helicase. Given the similarity of the mean track duration estimates to the photobleaching time using 1s interval, we could not calculate bound time estimates for these subunits under these conditions.

**Figure 3.**
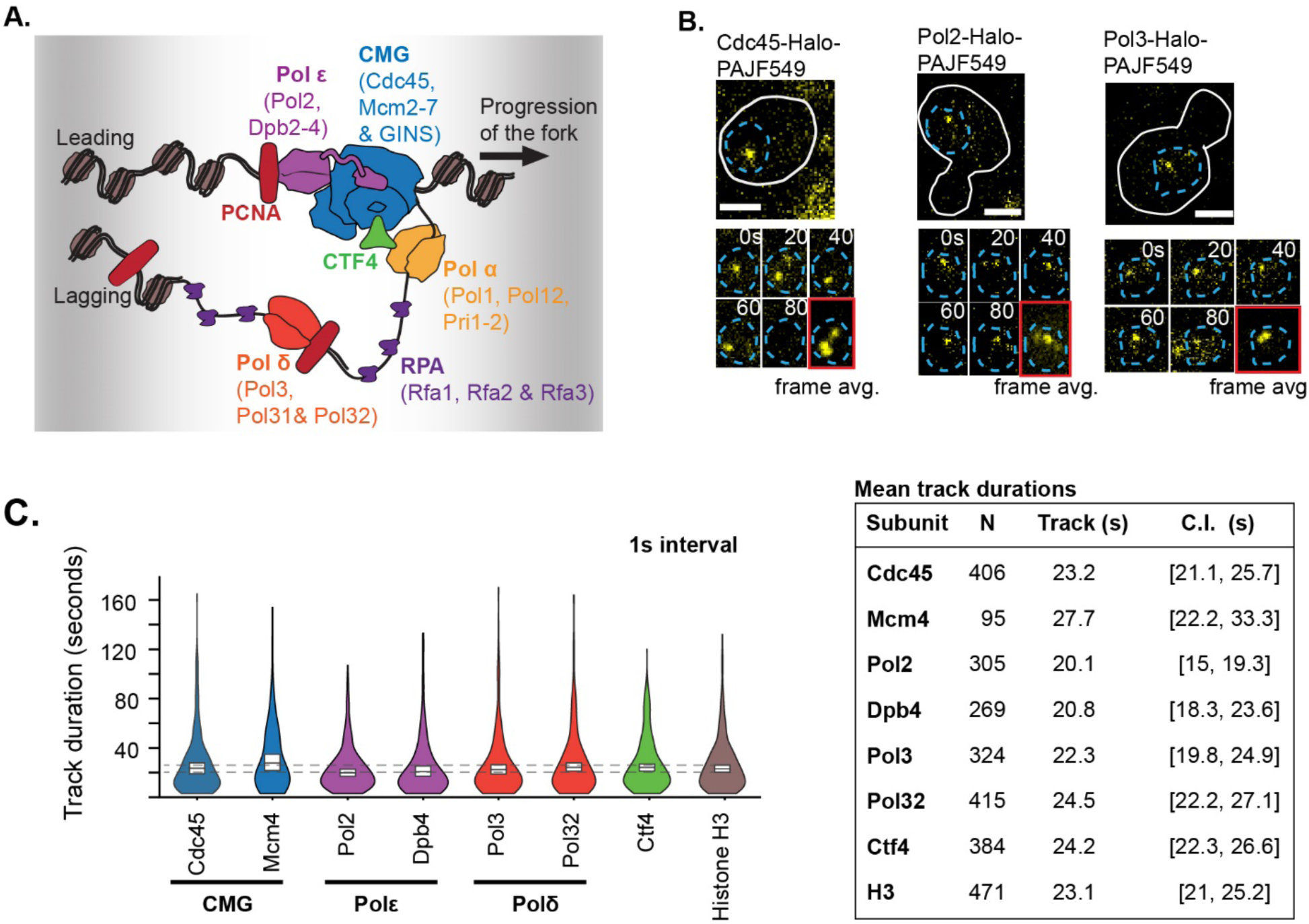
Binding kinetics of replisome subunits. (**A**) Diagram of the architecture of the replisome in *S. cerevisiae*. (**B**) Images for long-lived single-molecule fluorescent foci in strains carrying replisome fusions to Halo-PAJF549. Images on the top show the entire cells. Images on the bottom show a magnified view of the nuclei and show individual timepoints or a frame average of the experiment over 80 seconds. Scale bar 2 μm. (**C**) Distribution of track durations (spot lifetimes) for different replisome subunits imaged at intervals of 1 second (left). White boxes represent the boundaries of confidence intervals of the mean (represented as the line in the box). Dashed lines, representing the confidence intervals for Histone H3, are drawn for comparison. Colours correspond to the diagram shown in A. At least two duplicates were done for each subunit except for Mcm4, for which data comes from a single experiment. Sample sizes and number of repeats are described in Table S7. (Right) Table summary of estimation of mean track durations, after combining data from individual experiments, for different subunits collected with 1s time intervals.

We corroborated and extended the results above by doing similar experiments using longer time intervals between acquisitions. To reduce the likelihood of chromatin-bound molecules moving out of the focal plane, we imaged three and four different planes separated by 500nm, for 8s and 20s intervals, respectively, and performed maximum intensity projections (MIP) on the resulting z-stacks (Figure 4A). These time intervals allowed us to observe some separation between the mean track duration estimates and the photobleaching time – with the exception of Cdc45 –, thus allowing us to estimate the bound time (Table S7). These experiments suggest that the CMG helicase, Ctf4, Pol δ, and Pol ε, remain bound to chromatin for an average time that of at least 5 minutes (Figure 4C). Note that the similarity to the bleaching control results in a high uncertainty in our estimates. Furthermore, given that our measurements originate from replisomes at varying stages between initiation and termination, the real residence times may be longer than our estimates. Synchronization of cells would not solve this problem, as replisomes would still be activated at different times and would synthesize different lengths of DNA. To put these numbers into perspective, the average segment replicated by an individual replisome, defined as half the length of a replicon, is 18.5kb (Liachko et al., 2010). At an average replication rate of 1.6 kbp/min (Sekedat et al., 2010), the time for synthesizing this segment is ∼11 minutes. Our estimates are of a similar magnitude as the time required for completion of synthesis, which suggests few or no turnover events of the DNA polymerases from initiation to termination.

**Figure 4.**
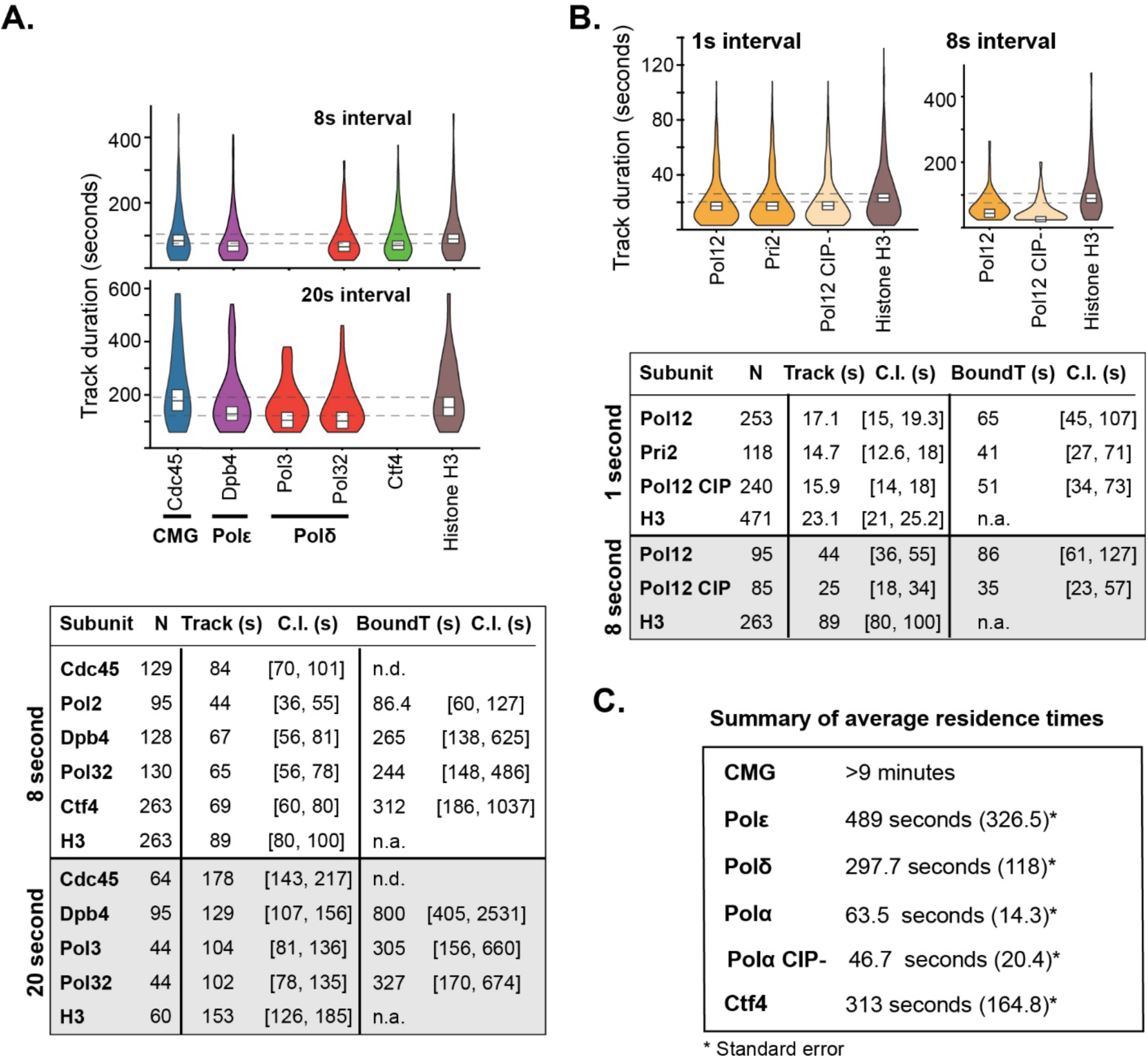
Estimation of bound times with longer time intervals. (**A**)(Top) Distribution of track durations for the same subunits as in Figure 3C using intervals between pictures of 8 or 20 seconds. (Bottom) Table summary of estimation of mean track durations and bound times for individual subunits. We could only calculate the bound time for cases where there was difference in the photobleaching time and track duration time, which is why we could obtain an estimate for Cdc45. Data for all subunits was obtained from at least two independent experiments. (**B**) (Top) Distribution of track durations for subunits for Pol α and comparison with histone H3, using 1s and 8s time intervals. Histogram for histone H3 is shown for these same distributions so that the difference with histone H3 can be appreciated. (Bottom) Table summary of the estimations of mean track durations and bound times for subunits of Pol α. Data for all subunits was obtained from at least two independent experiments. (**C**) Summary of the residence times obtained. For Cdc45, we placed a lower limit on its residence time (Materials and Methods, Figure S13). For the rest of the components, the estimates represent weighted averages from residence time estimates using equation 2 (Materials and Methods), that we could obtain at different time intervals and different subunits of a component (Tables S7).

Pol α was the only subcomplex that exhibited a more dynamic behaviour among the polymerases. Using 1-second intervals between consecutive pictures, both Pol12-Halo and Pri2-Halo had track durations shorter than the bleach time (Figure 4B). This observation was corroborated by results using 8-second intervals for Pol12-Halo. We also found no evidence supporting multiple binding behaviours, based on statistical tests that support the use of a single-exponential over a two-exponential model (Table S3). A weighted average of the two subunits and intervals results in an estimated residence time of 63.5 seconds, indicating that Pol α can perform multiple cycles of priming before unbinding (Figure 4C). To test if the binding kinetics of Pol α reflect a dynamic binding to the stably-bound Ctf4, we repeated the experiments of Pol12-Halo in a mutant strain of Pol1 lacking the Ctf4-interacting peptide (CIP), termed Pol12-Halo CIP-(Figure 4B). Surprisingly, while we detected a slightly lower residence time in the CIP-mutant when compared to the wt strain, the two values are still similar and within error of our estimates, suggesting that the interaction with Ctf4 may play only a minor role in retaining Pol α at the replication fork (Figure 4C). This is consistent with an *in vitro* study suggesting that Ctf4 does not seem to have an effect on the non-distributive behaviour of Pol α at the replisome in the context of chromatin, and an *in vivo* study that DNA synthesis is unperturbed and Pol α still remains associated to the replisome (albeit it to a reduced degree) when the Ctf4-Pol α interaction is abolished, suggesting a Ctf4-independent link of Pol α to the replisome(Evrin et al., 2018; Kurat et al., 2017). Ctf4’s role put aside, these results show that Pol α is the most dynamic of the three replicative polymerases.

### Validation of the single-molecule approach by FRAP

Given that some of the single-molecule microscopy results were unexpected, we sought to validate our approach using an independent technique. Fluorescence recovery after photobleaching (FRAP) was chosen as is commonly used to study protein dynamics in live cells, and we have used it before to study the dynamics of replisome components in *E*.*coli* (Beattie et al., 2017; Mueller et al., 2010). To do it, we generated strains carrying mNG C-terminal fluorescent fusions for multiple replisome subunits. The rationale of these experiments was to measure the fluorescence recovery of the chromatin-bound protein fraction for each replisome subunit studied, from which we could extract protein residence times. To strengthen our conclusions, we also applied the complementary technique, fluorescence loss in photobleaching (FLIP).

Our results with FRAP are consistent with those from single-molecule microscopy (Figure 5). Both Mcm4 and Ctf4 showed slow turnover rates that did not reach the maximum possible recovery over the length of our experiments (defined by the average intensity in the nucleus after the bleaching step) (Figure 5A-B). In the case of Mcm4-mNG, we estimated a single time-constant of 62 seconds, but the significance of this number is unclear since the curve did not get close to the maximum recovery and is therefore possibly reporting on nuclear and chromatin movement. In the case of Ctf4-mNG, we detected two time-constants, one of 7 seconds – likely representing the diffusive pool – and a second one estimated at 309 seconds – likely representing chromatin-bound fraction – we obtained lower estimates of 5.4 and 84 seconds for FLIP, respectively (Figure 5C). These estimates had a high uncertainty reflecting a small sample size and time-constants longer than our experiments. In contrast, Rfa1-mNG showed full recovery at a fast rate, with time-constants of 4.2 and 5.6 seconds for FRAP and FLIP, respectively (Figure 5A-C). In addition, we also applied this technique to PCNA-mNG, which was not previously characterized by single-molecule microscopy, and obtained residence times of 32 and 51 seconds for FRAP and FLIP, respectively (Figure 5A-C).

**Figure 5.**
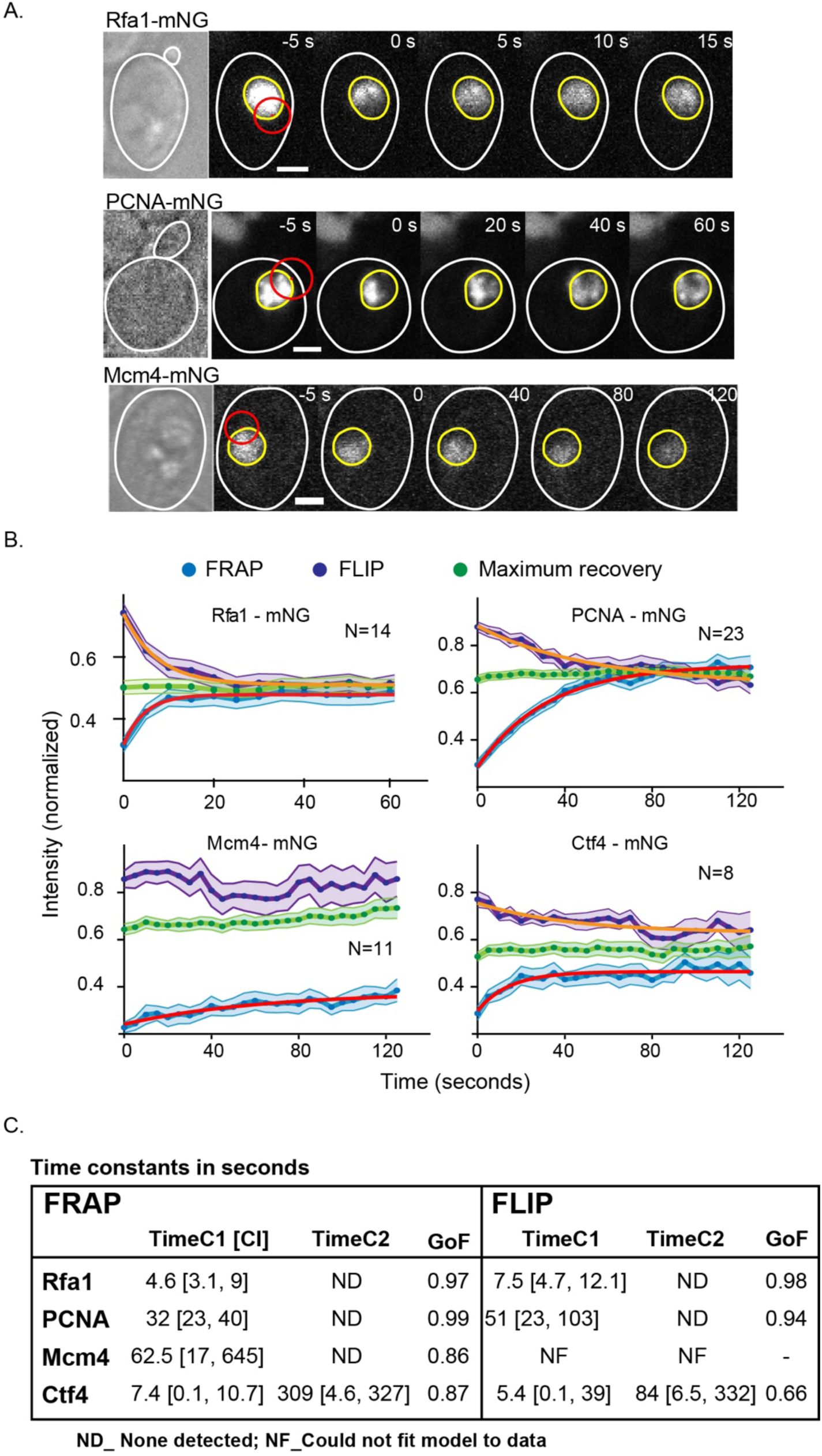
FRAP results of replisome subunits. (**A**) Representative images showing bleaching, and fluorescence recovery for Rfa1-mNG, PCNA-mNG and Mcm4-mNG. The 0-seconds time point represents the time at which selective photobleahing occurred. The red circle shows the point estimated bleached area. Scale bars represent 2 μm. (**B**) Plots showing the average recovery curves and their fitting. Standard errors of average points are shown in shaded colours. The maximum recovery, calculated as the average intensity of the nucleus after photobleaching, is shown in green. Model fitting of FRAP and FLIP curves are shown as red or orange lines, respectively. (**C**) Table summarizing the results after fitting the data to a single or double exponential model.

Our FRAP experiments were less adequate for the study of nuclear events in yeast than single-molecule microscopy. Multiple constraints, inherent to FRAP or particular to the study of the yeast’s nucleus, limited the sample size and increased the uncertainty in our estimates (discussed in Supplemental Text). In addition, FRAP likely requires a high chromatin-bound to diffusive ratio to determine residence times. In the case of DNA polymerases, over 90% of their copies are expected to be diffusing during most of S phase – based on their reported copy numbers and the maximum estimated number of replisomes at the peak of S phase (Ho et al., 2018; Saner et al., 2013). We think this was the reason why we detected no chromatin-bound fraction and could not obtain sensible estimates in our experiments for Pol3-Halo, Pol2-Halo, and Pol12-Halo (Figure S14). Dual labeling of these polymerase fusions and PCNA-mNG allowed us to verify that cells were in S phase and that FRAP had worked (Figure S14A, Pol3’s example). Despite not being capable of corroborating our estimated residence times of the DNA polymerases, overall our results with FRAP validated our single-molecule approach.

### Pol δ is likely linked to the rest of the replisome

Four different models can explain the observed stable-binding of Pol δ: first, stable binding after completion of Okazaki fragments; second, usage at both leading and lagging strands; third, efficient recycling and quick rebinding in replisome-dense nuclear regions; fourth, tight binding to the replication fork. The first of them, slow recycling after completion of the Okazaki fragments possibly through binding to PCNA, is incompatible with the estimated copy numbers of the Pol δ subunits. Considering that at the peak of S phase there are ∼300 replisomes per nucleus (Saner et al., 2013), and that the estimated copy number of Pol32 is ∼1600 (reportedly the least abundant subunit)(Ho et al., 2018), cells would only be able to undergo up to 5 cycles at the lagging strand, spanning about 30 seconds (165bp Okazaki fragments x 5 / 1.6kbp min-1 rate), before depleting the available Pol δ. In addition, our estimate for PCNA residence time is much shorter than that of Pol δ, indicating that this model is unlikely (Figure 5C). We acknowledge that the average Okazaki fragment size of 165bp, may be an underestimate as length distributions previously reported included longer fragments (Smith and Whitehouse, 2012). However, we believe our argument is valid even if we assume an average Okazaki fragment length 2-or 3-fold longer.

The second model, where Pol δ works at both strands, would potentially result in two different binding regimes: a residence time of several minutes as the DNA Pol δ elongates the leading strand, and a residence time of few seconds at the lagging strand comparable to the time needed to synthesize an Okazaki fragment. However, statistical tests designed to search for multiple behaviours showed no evidence for the presence of two regimes in our data (Figure 6A; Table S3).

**Figure 6.**
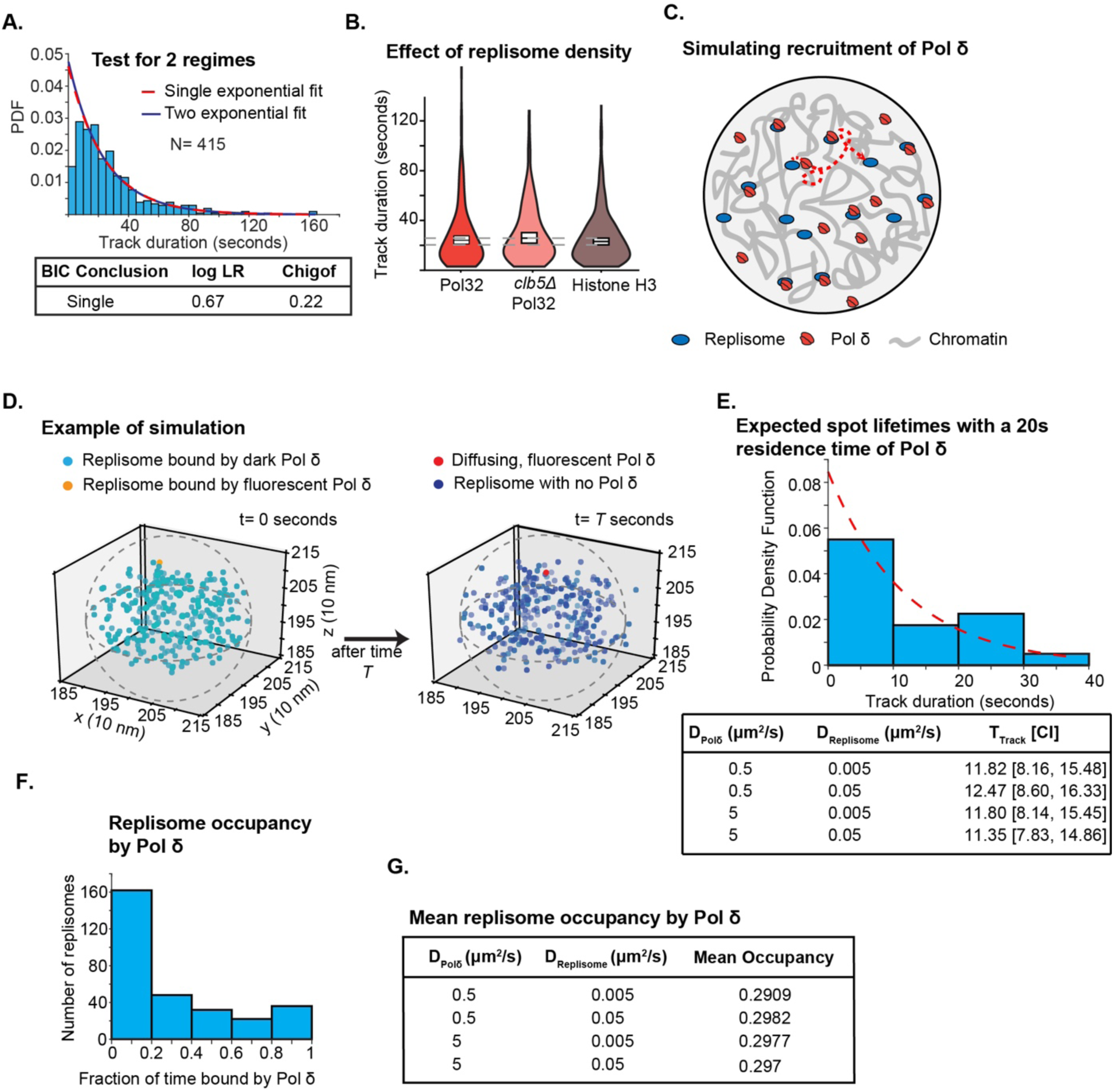
Testing for recycling of Pol δ. (**A**) Statistical testing of track durations obtained from Pol32 with 500ms interval, to check if a more complex model fits better than a single-exponential, indicating multiple binding behaviours at the replisome. The tests used were the loglikelihood ratio (LLR) test and the Bayesian information criterion (BIC) test, and results in the table show the p values from the test, where p<0.05 is deemed statistically significant (Table S3). (**B**) Comparison of track durations obtained of Pol32 in a *clb5Δ* background with a 1s interval, and from the histone H3 data. Data for Pol32 is identical to that in Figure 3C. (**C**) Diagram for recycling of Pol δ among multiple replisomes. (**D**) Diagram of general simulation setup zoomed into the sphere where all the replisomes were initially placed. Initially, 300 replisomes were placed inside a sphere that was 300nm in diameter to represent a diffraction-limited region, represented by the dotted line. Unbound, non-fluorescent copies of Pol δ are not shown for clarity. (**E**)Top – example of exponential fit on track durations. Bottom – track durations obtained from simulations, over a range of diffusion coefficients. The initial fraction of replisomes bound by Pol δ was 0.75, and the excess copy number of Pol δ was 1600. The estimated track duration was predicted to be 10.7 seconds, based on a 20s residence time and a 23s bleach time. (**F**) Representative histogram of the fraction of time replisomes had a bound Pol δ over the course of 200s. (**G**) Results for mean occupancy of replisomes (i.e. the average fraction of time replisomes had a bound Pol δ), over a range of diffusion coefficients. For both sets of simulations, we tested additional parameters summarized in Tables S4-S5.

The third model considers that Pol δ efficiently rebinds chromatin after completion of an Okazaki fragment. This idea is especially compelling due to the high density of replisomes at mid S phase. We expected that a lower replisome density would result in short dwell times if these were influenced by Pol δ recycling. To test this model, we artificially reduced the replisome density by using a strain carrying a deletion of *clb5*, which should result in the inactivation of about half of the origins of replication – specifically, inactivation of late origins (McCune et al., 2008)(Figure S3). The *clb5Δ* strain had a longer S-phase, bigger cells, and PCNA-mNG foci more distinguishable than wt (Figure S15), supporting the notion that fewer replisomes are active and that they take longer to duplicate the genome. However, our single-molecule estimates show no significant difference in the lifetimes of Pol32-Halo when comparing wt and *clb5Δ* cells, suggesting that fast rebinding cannot explain the long residence times observed (Figure 6B).

We also performed computer simulations of Pol δ binding in the context of high density of replisomes at mid S phase (Figure 6C-D). We initially placed 300 replisomes, each occupying a cubic space of 10nm side lengths, inside a sphere of diameter 300nm, corresponding to a diffraction-limited spot. This unnaturally high density was used to maximize the probability of recycling. Replisomes were permitted to move outside this sphere at rates reported previously for chromosomal loci, but were constrained in a larger sphere representing the nucleus without the nucleolus (diameter = 1.3µm). A fraction of 75% of all replisomes had a Pol δ bound to them initially – similar results were obtained with an initial fraction of 25% (Table S4). Assuming dissociation after Okazaki fragment completion, and factoring DNA synthesis as well as slowdown due to strand-displacement activity, we used a mean dwell time of 20 seconds for Pol δ, in our simulations. Unbound Pol δ molecules moved by Brownian motion until encountering a region occupied by a vacant replisome, upon which that Pol δ was considered to be bound to that replisome. We incorporated the bleaching time for 1-second imaging intervals (23 seconds), experimentally determined. To estimate track durations, we randomly assigned a Pol δ initially bound to a replisome to be in the fluorescent state and measured the number of time steps over which it remained fluorescent and replisome-bound (including binding to other replisomes). This simulation was repeated 40 times and fitted to an exponential model to estimate the mean track duration (Figure 6D, Video S2).

Our simulation results indicated no significant contribution of Pol δ rebinding under experimental and biologically sensible parameters. The estimates from simulations were consistent across the simulation parameters and ranged from 11.3-12.5 seconds (Figure 6E). This result clearly contrasted with experimental results using 1-second intervals showing track durations >22 seconds. Instead, the results were similar to a predicted mean track duration of 10.7 seconds for a model where there is no rebinding – based on a 20 second dwell time and a 23 second bleach time (equation 2, Materials and Methods). Furthermore, statistical tests favoured the use of a single-exponential over a two-exponential fit (Table S4). These two results suggested that bound fluorescent Pol δ molecules did not bind to a neighbouring replisome after dissociation. The result of our simulations also suggested that dissociation at each Okazaki fragment would result in a high proportion of replisomes not bound to Pol δ (Figure D). In a simulation over 200 seconds, and with all replisomes having a Pol δ bound to them, the mean occupancy of Pol δ at the replisome was only of ∼30%, but the distribution was highly heterogenous (Figure 6F-G, Table S5). In summary, simulation results not only argue against Pol δ recycling in our data but suggest that it would be incompatible with efficient DNA synthesis in replisomes distributed across the genome.

Based in the results above, we support the fourth model and propose that a single copy of Pol δ synthesizes multiple Okazaki fragments. This model would explain how the eukaryotic replisome achieves near identical rates of synthesis between the leading and lagging strand (discussed in (Kapadia and Reyes-Lamothe, 2019)).

## DISCUSSION

The classical model for the study of cellular replisomes is the bacterium *E. coli*. In *E. coli*’s replisome the evidence overwhelmingly points to a physical link between the replicative DNA polymerases and the helicase (Gao and McHenry, 2001; Kim et al., 1996). This physical connection permits the establishment of synchrony between helicase unwinding and polymerase activity (Kim et al., 1996; Sanders et al., 2010), which is presumably important to limit the generation of ssDNA during elongation of the strands. Here we provide evidence that supports a similar architecture for the eukaryotic replisome, where both Pol δ and Pol ε, which synthesize most of the genome’s DNA, are physically linked to the CMG helicase (Figure 7). Retaining a copy of Pol δ at the replication fork would facilitate its loading at PCNA and reduce the time that the ssDNA is exposed at the lagging strand. Physical interaction between helicase and polymerases may also provide a point of regulation in the rate of DNA unwinding by the CMG helicase after polymerase detachment, as is the case in bacteria (Dallmann et al., 2000; Kim et al., 1996). This physical link also implies the formation of single strand loops at the lagging strand as in the trombone model proposed by Alberts (Sinha et al., 1980).

**Figure 7.**
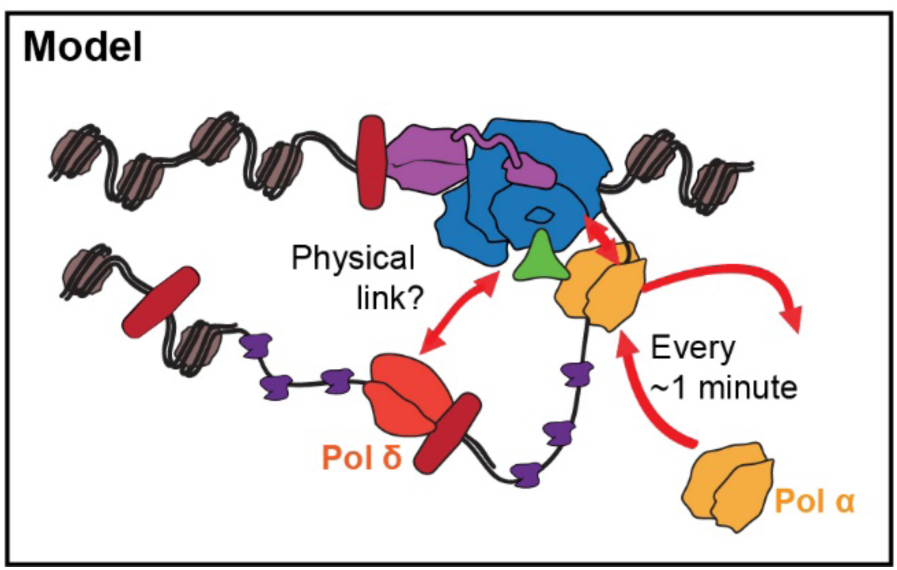
Model for the architecture of the eukaryotic replisome. The CMG helicase, Pol ε and Ctf4 bind tightly to chromatin and act processively. In addition, our data suggest the existence of a direct or indirect interaction between Pol δ and the stably bound CMG helicase. Pol δ acts processively, with a single copy of this polymerase typically synthesizing several – and possibly all in some cases – Okazaki fragments of an individual replisome from initiation to termination. Pol α interacts with the CMG independently of Ctf4 and turnovers at rates that permit it to synthesize multiple primers at every binding event. The function of the interaction between Pol α and Ctf4 at the replisome is unclear.

We hypothesize that a yet uncharacterized protein-protein interaction bridges Pol δ and the CMG helicase (Figure 7). It is possible that interaction with Pol α may help to retain Pol δ at the replisome (Huang et al., 1999); however, Pol α has a shorter residency time than Pol δ, indicating that there are likely other important interactions retaining Pol δ at the replisome. In addition, removal of Pol32 (the subunit of Pol δ that interacts with Pol α), showed only a small difference in the residence time of Pol δ *in vitro* (Lewis et al., 2020). In humans an interaction has been described between Pol δ and AND1, the orthologue of Ctf4 (Bermudez et al., 2010), which if present in budding yeast may contribute to retain Pol δ at the replication fork.

Recently, an *in vitro* single-molecule study on non-chromatinized DNA templates investigated the stability of eukaryotic polymerases, showing stable binding of polymerases to the replisome that agree with our results (Lewis et al., 2020). However, Pol α in their assay exhibited residence times of several minutes, leading to their proposal that it may participate in retaining Pol δ at the replisome. In contrast, our results suggest that while Pol α may be stable enough to perform multiple cycles of priming, it is not stable enough to explain the stability of Pol δ. Our results agree with a recent *in vivo* study suggesting Pol α may be dynamic as varying its concentration led to changes in the size of Okazaki fragments (Porcella et al., 2020). Furthermore, another study suggested that chromatin plays a role in retaining Pol α by showing differences in its stability on chromatinized and naked DNA templates(Kurat et al., 2017). The stability of Pol α and how it may influence the retention of Pol δ at the replisome will be of interest for future studies.

Residence times with minute-time scales of both the leading and lagging strand DNA polymerases in yeast contrast with the faster dynamics reported for the bacterial replisome (Beattie et al., 2017; Lewis et al., 2017; Liao et al., 2016), where DNA Pol III is exchanged every few seconds. This might be explained by differences in the accumulation of torsional stress near the replication fork, which has been implicated as a potential factor triggering unbinding of the polymerases in bacteria (Kurth et al., 2013). A ∼30 times slower DNA replication rate in budding yeast compared to *E. coli*, combined with about 10-fold shorter Okazaki fragments, may result in the accumulation of lower levels of torsional stress in budding yeast. Alternatively, strategies unique to eukaryotes may result in a similar transient binding of the DNA polymerases to DNA that do not require complete detachment from the CMG helicase. For example, despite of the stable binding of Pol ε to CMG, its flexible linker potentially allows it to unbind from DNA while remaining bound to the rest of the replisome (Zhou et al., 2017). Based on the resolution limits of our system, we cannot exclude the possibility of the polymerases being stable in the replisome, yet dynamic on DNA.

*S. cerevisiae* continues to be a powerful model organism for the understanding of basic eukaryotic cellular mechanisms due to its genetic tractability. However, the development of single-molecule imaging techniques, which have advanced the understanding and are widely applied in bacterial systems, have had limited use in this system due to higher background fluorescence and light scattering. Here we describe an approach to overcome these obstacles. We expect that the single-molecule protocols described here will facilitate its future use to study a wide range of questions in this organism.

## ACKNOWLEDGMENTS

We thank Tatiana Karpova (NCI) for providing the yeast strains carrying H3-Halo and pdr5Δ;Luke Lavis (Janelia Farms) for providing the Halo substrate coupled to PA-JF549 dye; and Karim Labib for the strain carrying the CIP-binding mutant of Pol1. We thank Thomas Gligoris and Stephan Uphoff for initial help working with HaloTag fusions. We also thank the students from the 2020 class of the course Biol518 in McGill University for help in proofreading and for comments on our manuscript. **Funding:** The experiments were done using equipment from the Integrated Quantitative Bioscience Initiative (IQBI), funded by CFI 9. This work was funded by the Natural Sciences and Engineering Research Council of Canada (NSERC# 435521-2013), the Canadian Institutes for Health Research (CIHR MOP# 142473), the Canada Foundation for Innovation (CFI# 228994), and the Canada Research Chairs program. **Author contributions:** NK, Conceptualization, Formal analysis, Investigation, Writing; ZWEH, Formal analysis, Investigation, Writing; HZ, Formal analysis, Investigation; AY, TRB, Investigation; RR-L, Conceptualization, Supervision, Funding acquisition, Writing. **Competing interests:** Authors declare no competing interests. **Data and materials availability:** All data, code, and materials used in this work are available by request.

## SUPPLEMENTAL MATERIALS

Supplemental Text

Figures S1 to S15

Tables S1 to S7

Videos S1 to S2

## MATERIALS AND METHODS

### Strains and constructions

Strains used in this study are all from a BY4741 background (with the exception of the CIP-mutant, see below) and are shown in Table S1. Plasmids used in this study are derivatives of pUC18 (ColE1 origin, Ampicillin resistance); pTB16 carries mNeonGreen and a downstream NatMX marker, while pSJW01 carries the HaloTag gene with a HygB marker. Both mNeonGreen and HaloTag genes encode an 8 amino acids linker at the 5’ end (sequence: GGTGACGGTGCTGGTTTAATTAAC). Plasmids were maintained in *E. coli* DH5α and were extracted by growing in LB with 100 μg/ml ampicillin then using the Presto Mini Plasmid Kit (Geneaid).

All PCRs were made using either Phusion or Q5 (NEB). PCR reactions were in a volume of 50 μl and included water, 3% DMSO, the reaction buffer, 2.5 mM of each dNTP, 0.2 μM of each primer, either 1ng of plasmid DNA (for insertions) or 1 μl of genomic DNA (for screening insertions), and 0.5 μl of polymerase.

Fluorescent fusions were made by PCR amplification from pTB16 or pSJW01 using the primers listed in Table S2. PCR products were transformed into wild-type diploid BY4743. A single colony was grown at 30°C in 5 ml yeast peptone dextrose (YPD) overnight, then diluted to a final OD600 of 0.1 in 10ml of YPD. Cells were taken at OD600=0.5-0.6 and centrifuged at 4000 rpm for 5 min. The pellet was washed twice with 25 ml of sterile deionized water, then once with 1ml of 100mM lithium acetate. Cells were then resuspended in 240 μl of 50% PEG, then 50 μl of salmon sperm DNA (thawed at 95°C for 5min, then incubated on ice for at least 10min), 50 μl of the PCR product and 36 μl of 1M lithium acetate were added in this order. The mixture was thoroughly mixed by pipeting and incubated on a rotator at 25°C for 45min, followed by heat shock at 42°C for 30min. Cells were pelleted in a microcentrifuge, washed in 500 μl of sterile water, then resuspended in 200 μl of YPD and plated on YPD agar. After growing at 30°C overnight, the cell lawn was replica-plated onto selective YPD agar, either with 100 μg/ml cloNAT (Werner) for mNeonGreen, or with 200 μg/ml Hygromycin B (Life Technologies) for HaloTag. Transformants were screened for the presence of an insert by PCR using the indicated primers in Table S2.

Confirmed clones were then sporulated by taking 750 μl of a YPD overnight cultures, washing 4 times with 1 ml sterile deionized water, then washing once with 1 ml of potassium acetate sporulation medium (Kac), and finally resuspending in 2 ml of KAc and incubating at 25°C with shaking. After 5 days the sporulating cultures were checked by microscopy for the appearance of numerous tetrads, then 750 μl was taken and washed 3 times in sterile water before final resuspension in 1ml water and storage at 4°C. For dissection, 45 μl of spores were treated with 5 μl of zymolase for 10 min, then tetrads were dissected on YPD plates to isolate haploids with the tagged fusion. Genomic DNA was isolated from the haploid by vortexing the cells in the presence of zirconia/silica beads, followed by phenol extraction and ethanol precipitation. The insertion site was once again amplified using the same screening primers as above, and the PCR product was sequenced to confirm that the tag and linker were both mutation-free.

The HaloTag haploids were combined with PCNA-mNeonGreen (from YTB31) and the *pdr5Δ::KanMX* deletion (from a haploid sporulated from YTK1414) by mating. To do this, 10 μl of water was spotted on a YPD plate, and colonies from the Mat a and Mat α haploids to be combined were mixed together into the water drop and incubated at 30°C overnight. Cells were then restreaked on an auxotrophic -Met-Lys plate on which only the mated diploid could grow. Diploids resulting from a mating were dissected as above. PCNA-mNeonGreen (from YTB31) and the *pdr5Δ::KanMX* deletion (from a haploid sporulated from YTK1414) were combined by mating and the resulting diploid was dissected to isolate strain ZEY098, a haploid with both PCNA-mNeonGreen and *pdr5Δ::KanMX*. ZEY098 was then mated with a haploid of the opposite mating type dissected from each of the HaloTag transformants to create haploids with all three markers (HaloTag, PCNA-mNeonGreen and *pdr5Δ*) that were then used for imaging and control experiments. These haploids are all listed in Table S1.

The CIP-strain YCE449 is from a W303 background and carries a *pol1-4A* allele, where four amino acids are substituted with alanine at D141, D142, L144 and F147 (Simon et al., 2014). Pol12-Halo and CIP-were combined by mating YJL10 with YCE449, then sporulating and dissecting the resulting diploid to get YAJ05.

### Western blot

To prepare crude cell lysates, YPD cultures were grown to exponential phase (OD600=0.5) and fixed in 10% TCA. The pellet was washed in cold acetone then in Beating Buffer (8 M urea, 50 mM ammonium bicarbonate, 5 mM EDTA), then resuspended in Beating Buffer with glass beads and vortexed at 4°C for 5min. The bottom of the tube was pierced with a needle, placed inside another tube and the cell debris and buffer was centrifuged into the clean tube. This was centrifuged again and the supernatant was collected. Protein concentration was determined using a Bradford assay.

SDS-PAGE was performed using 4-20% Mini-Protean TGX precast gels (Biorad). Lysates were prepared in Laemmli sample buffer with β-mercaptoethanol. Gels were run in Tris-Glycine-SDS at 100V for 2-3h. Proteins were transferred onto a nitrocellulose membrane using Biorad’s Trans-Blot Turbo system and transfer packs running the MixedMW preset program. The membrane was incubated in Blocking Buffer (5% milk and 3% BSA in TBS-Tween) for 1h. The membrane was probed with α-Halotag mouse monoclonal antibody (Promega, diluted 1:1000 in Blocking Buffer), washed 3 times in TBS-Tween, then probed with goat α-mouse HRP-conjugated secondary antibody (Promega, diluted 1:10000 in Blocking Buffer) and finally washed 3 times in TBS-Tween. The membrane was treated with Clarity Western ECL substrate (Biorad) and exposed to autoradiography film (Diamed).

### Flow cytometry

Cells were prepared for flow cytometry by growing cultures in YPD to exponential phase (OD600=0.5) and fixing them in 70% ethanol at 4°C. Cells were washed in Tris-Cl and incubated with RNAse A at 42°C for 3h, then with Proteinase K at 50°C for 30min. Cells were centrifuged and the pellet was resuspended in Tris-Cl and stained with propidium iodide (8 μg/ml). Samples were run on a FACSCalibur (Becton Dickinson) using the following filters and detector settings: FSC E01; SSC 396V and 4.61 gain; FL2 730V and 4.10 gain. Cytometer was calibrated using parental haploid (BY4741) and diploid (BY4743) asynchronous exponential cultures.

### Microscopy

A single colony from a YPD plate was placed in 5mL synthetic complete (SC) medium and grown with shaking at 30oC for ∼5-6 hours. This culture was diluted by transferring ∼50uL into 5mL of fresh SC and grown overnight at 30 degrees. The overnight culture was diluted to 0.15 the next day and grown until the optical density (OD) reached 0.30. 1mL of this culture was spun down for 1 min at 4000RPM, and the pellet was resuspended in 500uL of fresh SC. Janelia Farms photoactivatable 549 (JF-PA549) was added to the 500uL culture for a final dye concentration of 50nM, except for YTK1434-Halo (Histone H3), where a concentration of 10nM was used to compensate for the higher copy number. This culture was placed in a thermomixer at 30oC and 500RPM for 40 minutes. After incubation, 3 wash cycles using fresh SC were done to wash away unbound dye. After the final wash step, the pellet was resuspended in 50uL of SC, and 3uL of the culture was placed on an agarose pad consisting of SC and Optiprep (Sigma), within a Gene Frame (Thermo Scientific). The pad was made by taking a 2% agarose Optiprep mixture (0.02g in 1mL Optiprep) - that was heated to 90 degrees - and mixing 500uL with 500uL 2xSC, resulting in a 1% agarose 30% Optiprep SC mixture. Approximately 140uL of this mixture was placed within the Gene Frame, with excess being removed with a KimWipe. Prior to imaging, we waited ∼15 minutes to let any unbound dye be released.

Coverslips were cleaned with the following steps: 1) Place in 2% VersaClean detergent solution overnight. 2) Wash with MilliQ water 3x. 3) Sonicate in acetone for 30 minutes. 4) Wash with MilliQ water 3x. 5) Place in methanol and flame coverslips using Bunsen burner. 6) Place in Plasma Etch plasma oven for 10 minutes.

Microscopy was done at 23 degrees, on a Leica DMi8 inverted microscope with a Roper Scientific iLasV2 (capable of ring total internal reflection fluorescence (TIRF)), and an Andor iXON Ultra EMCCD camera. An Andor ILE combiner was used, and the maximum power from the optical fiber was 100mW for the 405nm wavelength, and 150mW for the 488nm and 561nm wavelengths. The iLasV2 was configured to do ring highly inclined and laminated optical sheet (HILO), for selective illumination and single-molecule sensitivity. Metamorph was used to control acquisition. A Leica HCX PL APO 100x/1.47 oil immersion objective was used, with 100nm pixel size. Any z-stacks were doing using a PInano piezo Z controller.

Single-particle photoactivated localization microscopy (sptPALM) experiments were performed by activating molecules with low power (0.5-2% in software) 405nm light to photoactivate ∼1 molecule/cell, followed by stroboscopic, long-exposure (500ms) illumination with 561nm light (5% in software) to image primarily bound molecules. The time intervals used were 0.5s, 1s, 8s, and 20s, with cycles of activation every 40 timepoints for the 0.5s and 1s interval data, and every 15 and 10 timepoints, for the 8s and 20s interval data, respectively. For the 8s and 20s time intervals, a mini 561nm z-stack of 1um and 1.5μm was done, respectively, with 0.5μm step sizes, in order retain molecules in focus. Maximum intensity projections (MIP) of these stacks were used for subsequent tracking analysis. A brightfield image and a z-stack of 6μm (0.3μm step size) in the 488nm channel, was taken before and after each timelapse, to ensure normal cell health and to find cells undergoing S phase through the presence of PCNA-mNG foci.

### Tracking Analysis

#### Aside from tracking, all analysis was done using Matlab

Tracking was done using Trackmate (Tinevez et al., 2017). First, molecules were localized in each frame using a Laplacian of Gaussian (LoG) method, with an estimated diameter of 2.5 pixels. An intensity threshold was chosen that was slightly low, to still detect molecules that were moving out of the focal plane. Subsequent classification helped discard potential false-positive tracks from analysis that will be discussed later. After localization, tracks were formed using the Linear Assignment Problem algorithm by linking molecules in consecutive frames. The linking distance was chosen based on calculating the cumulative distribution function of the step sizes from the Histone H3 data, thus providing information on the step size of chromatin-bound proteins. Essentially, we varied the linking distance when analyzing the Histone H3 data until a noticeable plateau in the cumulative distribution function (CDF) was observed. We determined the linking distance as being the step size giving the 0.95 probability value in the CDF and did this for multiple time intervals: 3 pixels for 0.5s and 1s interval, 5 pixels, and 7 pixels for 8s and 20s time intervals, respectively. A gap frame – to allow for missed localization – of 1 was used the gap-linking distance was set to 2 pixels more than the linking distance for that time interval (e.g. 5 pixel gap-linking distance when using a 3 pixel linking distance). Linking also had cost of 0.3 for the “Quality” parameter to ensure that correct molecules were linked. We also allowed for track merging and splitting with a 5 pixel distance for all time intervals, in some cases where multiple molecules were activated within the region and they were near one another. Tracks with less than four localizations were discarded as they were unreliable. Furthermore, since we are using track merging and splitting, we can exploit this property to discard tracks that are found in noisy regions or with too many molecules active. In these cases, one would expect more spots localized in a track than predicted given a certain track duration (e.g. for a track duration of 20 frames, one would expect 21 spots, but may get 40). Therefore, we calculated a ratio of number of spots in track vs. number of spots estimated based on track duration and used a cut-off of 1.5 to remove tracks with too many spots localized (Figure S7).

To isolate tracks found in the nuclei of cells undergoing S phase, we used the PCNA-mNG z-stacks, as PCNA is active during S phase, thus resulting in fluorescent foci (Kitamura et al., 2006). First, we performed a Smooth Manifold Extraction (SME) (Shihavuddin et al., 2017) on the z-stack, as an alternative to MIP, to distinguish more clearly PCNA-mNG foci. We then generated a binary mask of this image, resulting in all the nuclei regions have intensity values of 1, and zero elsewhere. To isolate S phase nuclei, we used a threshold on the standard deviation of the intensities within the regions, as an indicator of heterogenous intensity caused by fluorescent foci. From the resulting binary mask, we isolated tracks found only in S phase nuclei by calculating the mean positions of tracks and determining if those positions land in the binary mask. For robust classification of tracks representing bound molecules, we employed a machine learning (ML) approach, using the output from Trackmate. First, we generated a training data set of ∼750 tracks from the Histone H3 data, taken with 0.5s interval (Table S8, data date:20180511). Given that we used long-exposure to blur out diffusing molecules, it is quite easy to detect bound molecules from the raw timelapses. Therefore, we manually classified tracks by assigning a value of 1 to tracks representing genuine bound molecules vs 0 for tracks representing diffusing molecules or noise. Once we had the classifications, we performed the learning procedure using the algorithm, Random Forest ((Breiman, 2001)), to develop two classification models, referred to as Model 1 and Model 2, with out-of-bag errors (equivalent to estimated cross-validation error) of 0.10 and 0.05, respectively. Model 1 had the predictor variables from Trackmate: mean speed, max speed, min speed, and median speed. Model 2 had all those variables as well, but in addition, included max quality. Both models were built with 6000 trees during the learning procedure, with following hyperparameters: InBagFraction (fraction of training data set given to each tree) = 0.50, MinLeafSize (minimum leaf size) = 50, NumPredictorstoSample (number of predictors to sample at each node) = 2. We then could output the predictor variables from tracks derived from timelapses of replisomal proteins, and input them into the models, and they would classify tracks as being bound.

The motive for building two classification models was to scale the predictor variables accordingly based on different time intervals and quality of data (e.g. slightly different laser intensities), compared to the training data set. First, we calculated the mean of the mean speed and the mean of the max quality from the Histone H3 training data set. Next, we performed a Gaussian Mixture Model (GMM) fit, with two components, on the log-transformed mean speed values of a given data set. We then took the mean of the first Gaussian component, likely representing bound molecules, and with the mean of mean speed values obtained from the training data set, we scaled all 4 speed values accordingly. For example, if the mean of mean speed from the training data set was 1.71 and the mean of the first Gaussian component was 2.5 (assuming a longer time interval), then the scaling factor would be 1.71/2.5 = 0.68. We then performed classification with Model 1.

From the tracks extracted with Model classification, we again performed a GMM fit, but this time on the log-transformed max quality values (Figure S6).

We then took the mean of the second Gaussian component (likely representing genuine molecules), and used the mean of the max quality value from the training data, to scale the quality values in a similar manner as described before. This would help ensure that despite slight differences in the quality of data compared to the training data set, tracks would still be robustly classified. After scaling, we input the tracks into Model 2, as the final classification step.

We performed a GMM fit, with the Bayesian Information Criterion (BIC) test incorporated to prevent overfitting, on the mean intensities of tracks followed by clustering, to isolate tracks representing single-molecules and not multiple ones (Figure S8). Also, given that we use very low dye concentrations, thus <<100% dye labelling, and the very low laser power for activation, it is unlikely that we are activating multiple molecules at once.

We then fit the track durations of the remaining tracks with a truncated exponential model, to compensate for discarding short duration tracks, using MLE through Matlab’s “mle” function, to calculate the mean track duration:

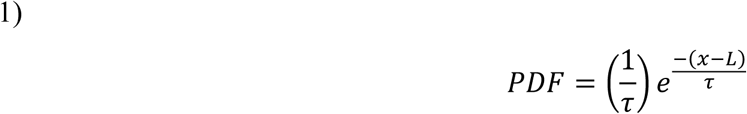

where τ is the mean track duration, and *L* is the truncation point (Balakrishnan and Basu, 1995). The 95% confidence intervals were calculated by bootstrapping 1000 samples. Bound times were calculated using the following equation:

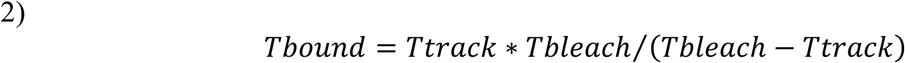

To calculate the errors on the estimate, we performed bootstrap sampling on the track durations from the combined data set (when multiple data sets of the same time interval were taken) to the following equation:

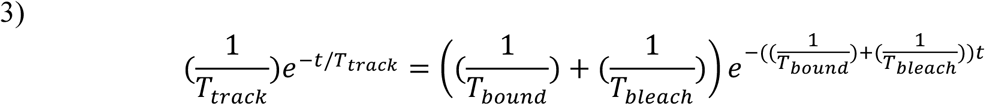

With 10% variation allowed for the *T*_.bleach_ estimate, in order to obtain biologically sensible results.

For Cdc45, since its track durations overlapped strongly with bleaching duration, we could not calculate a reliable estimate, so we estimated a minimum bound time based on the bleach duration with 8s time interval (Figure S13). Using equation 2, we calculated track durations by varying the bound time with a fixed bleaching time. The minimum bound time we chose was the one that gave us a track duration time equal to 1 second lower than the lower bound of the CI from the estimated bleaching time.

To check for two-exponential mixtures, the track durations were fit with the following two-exponential model:

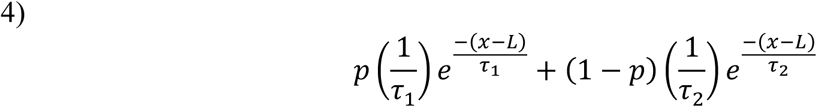

where τ_1_ = (*Tbleach + Tboundalpha*)/(*Tbleach* \**Tboundalpha*), τ_2_= (*Tbleach + Tboundbeta*)/(*Tbleach * Tboundbeta*), *p* is the mixture proportion, and *L* is the truncation point.

The lower and upper bounds on the two binding timescales were 0.1s and 900s, respectively, while allowing for a 20% variation in the bleaching estimate.

To check for overfitting and to identify whether the two-exponential model significantly fit the data better, we used the BIC test and the Loglikelihood ratio (LLR) test, as described in (Beattie et al., 2017). We looked for cases when the two-exponential model estimates did not simply return the lower and/or upper bounds as this would indicate that no physically sensible solution was found. We also performed a chi-square goodness of fit test, under the null hypothesis that the data comes from a single-exponential distribution.

### Simulations of Pol δ

#### Simulations were performed using Python 3.7

A 3D array of 400 × 400 x 400, was generated, with each element corresponding to a 10nm x 10nm x 10nm region. The yeast nucleus has a diameter of ∼ 1.5um, but given that the nucleolus encompasses roughly 1/3 of the nuclear volume, we modelled the confinement volume as being a sphere of diameter 1.3um (Taddei and Gasser, 2012). Inside the nucleus was another sphere – termed the replisome sphere – that was 300nm in diameter, to resemble a diffraction-limited region. Both spheres had their centers placed at position (200, 200, 200) of the 3D array. Replisomes were placed randomly within the replisome sphere but are subsequently allowed to move throughout the nucleus. A fraction of Pol δ were bound to replisomes while the unbound fraction (referred to as excess copy numbers) were placed throughout the nucleus. One of the bound Pol δ molecules was assigned to be fluorescent, in order to test for any rebinding behaviour of a fluorescent molecule. To simulate diffusive behaviour of the unbound fraction, we calculated the step size in each direction, by sampling from a normal distribution of mean 0, and a standard deviation of 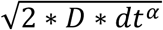, where *dt* is the time lag (0.05 seconds), α is the diffusion exponent (equal to 1 for diffusing Pol δ, and 0.5 for replisomes, representing chromosomal loci movement (Hajjoul et al., 2013)), and *D* is the diffusion coefficient. If the step size caused the molecule to go out of the nucleus, the step size was resampled until it remained in the nucleus. Movement of replisomes based on genomic loci movement, was assumed to be subdiffusive as explained by the Rouse model (Hajjoul et al., 2013). To represent the 200 seconds of acquisition time for the 1 second time interval data, we performed 200s/(0.05s/time step) = 4000 time steps. The residence time of Pol δ – assuming one copy per Okazaki fragment – was estimated to be 20s, based on the size of Okazaki fragments measured *in vivo* (∼160bp) (Jackson et al., 2012), the amount of RNA/DNA removed by Pol δ through strand displacement activity, and the rate of synthesis of Pol δ under physiological conditions. The rate of synthesis and strand displacement activity has been reported to be ∼50bp/s and 5bp/s, respectively at 30 degrees Celsius (Stodola and Burgers, 2016), so we took half those rates given we performed experiments at room temperature, given the replisome speed at room temperature is roughly half the speed at 30 degrees Celsius (Raghuraman et al., 2001; Sekedat et al., 2010). We assumed that 130bp are synthesized at 25bp/s (∼5s) while 30bp representing an DNA/RNA primer synthesized by Pol α (Stodola and Burgers, 2016), is removed through strand displacement activity at a rate of 2.5bp/s (12s), resulting in a predicted residence time for Pol δ of 17s. We used 20s in the simulation to allow for delays in the engagement with PCNA. The excess copy number of Pol δ used in the simulation was 1600 copies, which is the estimate of the lowest abundant subunit of Pol δ, Pol32 (Ho et al., 2018).

Two sets of transition probabilities were setup; one for the fluorescence state kinetics (e.g. photobleaching), with two states (fluorescent vs. bleaching) and the other for the molecule kinetics (e.g. unbinding), with two states (bound vs. unbound) since these were assumed to be independent from one another. Binding of Pol δ to a replisome was assumed to happen when the molecule entered a region occupied by a replisome and bound molecules would move along with their respective replisomes until dissociation. The next state of the molecule after a time step, was calculated by a sampling from a multinomial distribution with proportions given by the appropriate transition probabilities.

To determine rebinding of a fluorescent Pol δ, we calculated the overall time it spent in the fluorescent, bound state, regardless of the specific replisome it was bound to. This gives us a track duration time for one molecule within a nucleus. A fraction of 0.75 of the replisomes had a Pol δ bound at the start of the simulation. We also performed with the simulation with a fraction 0.25 with no significant effect (data not shown). We repeated this simulation 40 times (representing 40 nuclei sampled) under the same parameter set, in order to obtain a distribution of track duration times, which we can fit to using MLE, to extract the average track duration. Based on the bound time we input (20 seconds), and the measured photobleaching time with 1s interval (23 seconds), we can calculate the predicted track duration using equation 2, representing a model where there is no rebinding, and compare it to the estimated average track duration from the simulations.

For the simulations regarding the occupancy of replisomes by Pol δ, they were identical to the ones described above, except we only performed one simulation per parameter set, and all replisomes were bound by a Pol δ initially. We then calculated for each replisome, the fraction of time it was bound by a Pol δ, across a 200s simulation duration.

### Diffusion Coefficient

Diffusion coefficients were estimated from individual tracks by calculated the slope of the time lag (τ) vs mean squared displacement (MSD) curve. The equation used was modified from (Michalet and Berglund, 2012):

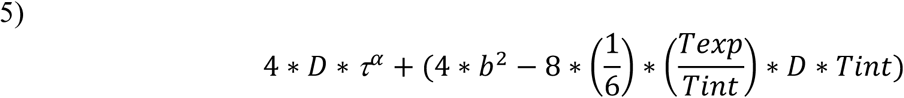

Where *D* is the apparent diffusion coefficient, α is the anomalous diffusion constant, *b* is the static localization error, *Texp* is the camera exposure time, and *Tint* is the time interval. We used this equation to correct for dynamic localization error due to the molecule moving within the exposure time.

To gain a better estimate of the diffusion coefficient, we performed a weighted least squares fit using the MSD values averaged over the tracks. We did not include tracks with a goodness of fit (gof) score of less than 0.70 from the previous step. We also included lower and upper bounds on the estimates to ensure physically sensible estimates, and in cases where on the estimates was the lower or upper bound value, we discarded data points at higher τ values. This is a known issue given there are fewer data points in the calculation of MSD at the higher τ values leading to more noisier traces.

### S phase vs nonS phase comparison of tracks

To determine if molecules found in S phase nuclei are also found in non S phase, we first segmented and selected for non S phase nuclei by selecting for those that had a homogenous intensity of PCNA-mNG (i.e. a low standard deviation of intensity within the nucleus). Then we ran our ML approach on tracks found within these regions. We scaled the speed variables and quality variables by using the same scaling calculated with the tracks found in S phase nuclei. After the ML classification, we identified which nuclei the tracks belonged to and which nuclei did not contain any classified tracks to generate a histogram of tracks/nucleus for both S phase and non S phase nuclei.

### No HaloTag control

Experiments were done similarly to those performed on replisomal subunits, except we used the strain ZEY098, which did not contain the HaloTag fusion, and thus would allow us to determine non specific binding of the HaloTag ligand, JF PA549. The dye concentration used was 50nM and images were acquired with 500ms with no time interval. S phase nuclei were identified based on PCNA-mNG segmented as mentioned previously. For ML classification, we used scaling factors calculated from replisomal subunits collected on the same day, to identify tracks resembling the tracks found within S phase nuclei of replisomal subunits fused to HaloTag.

### Cell cycle studies of Halo-tagged replisome proteins

For the study of the relation between spot appearance and the cell cycle, cultures were grown and stained with the PAJF549 dye and slides were prepared exactly as described above for the single-molecule experiments. All strains had PCNA tagged with mNeonGreen (POL30-mNeonGreen), a *pdr5* deletion (Δ*pdr5*::KanMX) and HaloTag fused to either Pol12 (YJL10), Pol2 (YJL02) or Pol32 (YJL11).

Imaging was performed at room temperature on an inverted Olympus IX83 microscope using a 100x oil objective lens (Olympus Plan Apo 100X NA 1.40 oil). Images were captured using an Andor Zyla 4.2 sCMOS camera. Excitation lasers originated from an iChrome Multi-Laser Engine (Toptica Photonics) and triggered through a U-RTCE real-time controller (Olympus). Experiments were done using HiLo illumination setup from a single-line cell^TIRF illuminator (Olympus). Olympus CellSens 2.1 imaging software was used to control both the microscope and lasers.

Image capture was performed as follows. First, 561nm laser at 12% power was used for 10s continuously to bleach pre-activated molecules (this helps reduce cellular background). This was followed by a single activation event of the Halo dye with 405nm laser at 15% power for 200ms (under our usual setup on this microscope, we use the same settings and 20ms to activate on average a single molecule per cell. Using 200ms ensures overactivation and therefore allows detection of multiple molecules per cell while minimizing bleaching and phototoxicity). Timelapse imaging was then immediately started with the following sequence of excitation and exposure times: bright-field imaging for 20ms, then 488nm at 14% power for 1s (for PCNA-mNeonGreen), then 561nm at 12% power for 1s (for Halo-tagged polymerase). This sequence was repeated a total of 30 times with a 2min time interval between frames.

Images were analyzed using custom scripts, and each frame of the timelapse was treated as a separate image. Using the green channel, cells were segmented into S phase or non-S phase depending on the presence of distinct PCNA foci or more diffuse nuclear fluorescence, respectively. Another script used the red channel to detect spots of polymerase single-molecules by fitting to candidate spots (local maxima), using an elliptical Gaussian function. These spots were then assigned to S or non-S nuclei based on the PCNA segmentation. A goodness of fit (gof) threshold was set to filter out low quality spots. The resulting data was used to plot the number of spots per nuclei for both and S and non-S phase cells.

To determine if the average number of spots per cell increased as cells got closer to mid-S phase, we used the results from the output as described in the previous paragraph, but on a cell-by-cell basis over time. We identified for individual cells the time frame at which the segmentation script detects a change in the categorization of a nucleus and set this as t0 for this nucleus, then we used the spot detection output to count the number of Halo spots for each cell before and after this t0. The average spot/cell was then plotted separately for cells entering from G1 to S phase, and cells exiting from S to G2.

To determine if the average number of spots per cell increased as cells got closer to mid-S phase, we used the results from the output as described in the previous paragraph, but on a cell-by-cell basis over time. We identified for individual cells the time frame at which the segmentation script detects a change in the categorization of a nucleus and set this as t0 for this nucleus, then we used the spot detection output to count the number of Halo spots for each cell before and after this t0. The average spot/cell was then plotted separately for cells entering from G1 to S phase, and cells exiting from S to G2.

### Fluorescence recovery after photobleaching (FRAP)

For FRAP experiments, the preparation of cultures and slides were done in the same way as single-molecule experiments. Imaging was done at room temperature, on a Leica DMi8 inverted microscope. A spinning disk imaging system (Andor Diskovery) was used, with a Roper 12 Scientific iLasV2, an Andor 13 iXON Ultra EMCCD camera and a Leica HCX PL APO 100x/1.47 oil immersion objective. The software Metamorph was used to acquire images.

A brightfield image was first acquired as a reference. Only S phase cells were selected for FRAP experiments. The selection was based on the bud appearance of the cells and the PCNA foci for strains carrying PCNA-mNeonGreen. FRAP was performed by using 405nm laser at 10–15% power, lasting around 300ms. The radius of the bleach region was around 1 µm. Given the large area bleached relative to the nuclear dimensions the selected bleaching region was centered at the edge of the nucleus, aiming to bleach half of the nucleus. Four pre-bleach images were acquired. After bleaching, 24 images were acquired. For mNeonGreen-tagged subunits, the image acquisition was done by using 200ms exposure at 5% 488nm power, with the time interval of 5s. For the strains with Halo-tagged proteins combined with PCNA-mNeonGreen, the images were acquired at 5% 488nm power and 15% 561nm power sequentially with the same exposure time and time interval as before.

### FRAP analysis

Images were analysed by using custom Matlab scripts. Since the bleached regions were often out of the targets, the filter step was first done to get rid of cells that did not bleached properly. The cells that underwent nuclear movement during imaging were also discarded. For selected cells, the sub-regions of bleached and unbleached region were manually defined. The nuclear regions of selected cells were defined based on their intensity using the custom script. The nuclear regions of three unbleached cells were also chosen as the bleaching control. A region not containing any cell was also selected background correction at each field of view. Intensity changes of all region of interests (ROIs) over the whole movie were obtained by using custom scripts. The FRAP, FLIP and maximum recovery curves were generated from the intensities of the bleached regions, unbleached regions and nuclear regions, respectively.

To correct for the differences in the initial intensity, acquisition bleaching and fluctuations in laser intensity, we normalized the intensity time traces. The corresponding average background value was first subtracted for each curve. Then the double normalization was applied by using the formula in (Koulouras et al., 2018).

Then the average FRAP curve was fitted both by single and double term exponential equations:

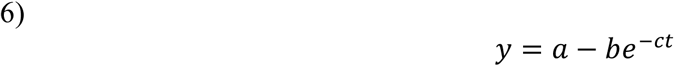

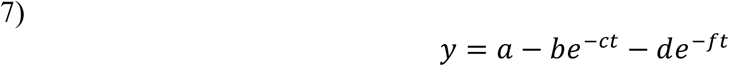

The average FLIP curve was also both fitted by single and double term exponential equations: 8)

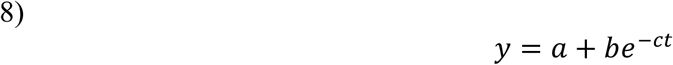

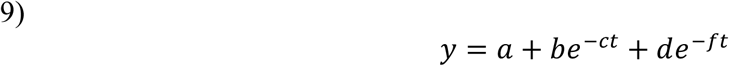

where *a* is the asymptote for recovery, *b* and *d* are the amplitude of recovery or loss, *c* and *f* are the rate of unbinding.

For the curve fitting, the non-linear least squares function in Matlab was executed. The upper boundary of *a* for FRAP curve fitting was set to the mean maximum recovery value plus 10%, while the lower boundary of a for FLIP curve fitting was set to the mean maximum recovery value minus 10%. The grid search of starting values was applied, resulting in many different fitting models. To determine the most suitable starting value, the adjusted R square value was used to assess the goodness of fit. The model with the highest goodness of fit was selected. To select between the single and double exponential fitting models for each curve, the Bayesian information criterion (BIC) was calculated. The model with lower BIC was selected. For curves that have similar BIC values, bootstrap sampling was applied. Single and double exponential models were fitted on each resampling dataset. Corresponding BIC values were then calculated. After 500 iterations, the model with the higher majority vote score was selected. The 95% confidence intervals (CI) of the estimated parameters were calculated by bootstrap sampling over 2000 samples.

The residence time were then calculated from the fitted results. The residence time equals to 1/*c* for both FRAP and FLIP single fit models, while equal to 1/*c* and 1/*f* for both FRAP and FLIP double fit models. The mobile fraction of the single fitted FRAP curve is:

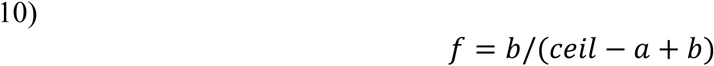

The mobile fraction of the single fitted FLIP curve is:

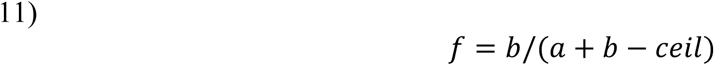

The two mobile fractions of the double fitted FRAP curve are:

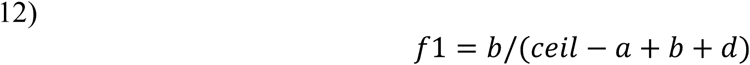

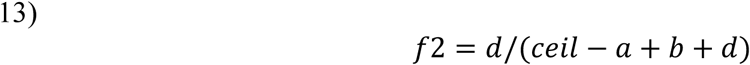

The two mobile fractions of the double fitted FLIP curve are:

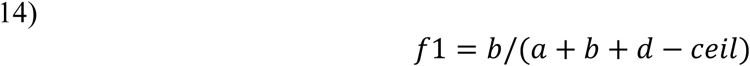

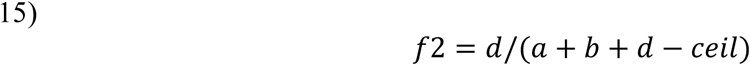

where *ceil* is the average value of the maximum recovery curve.

