## Supplemental Information for "Processive activity of the replicative DNA polymerases in the replisome of live eukaryotic cells"

### Supplemental Text

#### Optiprep viability

Iodixanol (Optiprep) was shown to be compatible with a variety of live specimens and to have no effect on their growth or viability (Boothe et al., 2017). Since none of the specimens tested had been yeast cells, we decided to verify that it did not affect growth and viability of *S. cerevisiae* under our specific experimental and imaging conditions.

In an initial attempt, we tried to perform growth curves on parental yeast strains with and without Optiprep. Due to the refractive index matching of Optiprep, the light absorption (Fig.S1A) and subsequently the OD measurements of cultures with the compound are lowered by an order of magnitude, which makes comparisons more difficult as the measurements are close to the lower detection limit of a spectrophotometer, which increases measurement error (Fig.S1B).

To alleviate this problem we approached growth measurement differently, by spotting YTB31 cells on Geneframes with SC-agarose pads, where the agarose is made in either water or in Optiprep (final concentration 30%). Cells were imaged in bright-field at 5min intervals over 15h at 22°C, with the Geneframe preventing evaporation and drying of the agarose pad. Each cell was then analyzed for cell cycle duration, by measuring the time between the appearance of a bud and the appearance of the next bud after division has occurred (Fig.S1C). The average of this time for all cells form multiple fields of view was used as a measure of doubling time. The doubling times with and without Optiprep were almost identical (125min and 126min, N=53 and N=31, respectively), which confirms that the presence of Optiprep in the agarose pad does not affect growth, and therefore is unlikely to affect DNA replication.

#### Growth and DNA replication in Halo-tagged strains

To ensure the presence of the Halotag at the C-terminus of replisome proteins, the mNeonGreen fusion to PCNA and the deletion of *pdr5* were not affecting DNA replication, we performed growth curves on the strains we used for imaging and compared this to the parental strain BY4741 as well as YTB31 (PCNA-mNeonGreen fusion alone). Cultures were grown in SC in the same way used for imaging (see Microscopy). OD was measured at 30min intervals over 9h and the slope of the resulting curve was used to calculate the doubling time of each strain. The average of two repeats is shown in the figure. The average doubling times of Halo-tagged Pol2 (YJL02), Pol3 (YJL24), Pol32 (YJL11), Mcm4 (YAY256), Cdc45 (ZEY158), Ctf4 (ZEY077), and the combined Pol12-Halo CIP- mutant (YAJ05) are all similar to those of BY4741 and YTB31 (Fig.S2), suggesting the genetic modifications in these strains do not affect growth and viability. In contrast, a strain where Dpb2 (another subunit of Pol ε) was tagged shows a higher doubling time than any of the other strains, indicative of potential growth problems. This strain was therefore not used for imaging. The histone H3-Halo strain YTK1434 (Ball et al., 2016) also showed significantly slower growth rate, though this should not affect our results since it was only used as a bleaching control.

Although we were confident the Halo tags did not affect growth, we analyzed the imaging strains by flow cytometry (see Methods) so we could be certain it was not causing more subtle problems with DNA replication that could affect our estimates of residency times without affecting growth rates. Strains with Halo-tagged Pol12 (YJL10), Pol2 (YJL02), Pol3 (YJL24), Pol32 (YJL11), Mcm4 (YAY256) and Cdc45 (ZEY158) all showed similar profiles to the parental BY4741, with a similar proportion of cells in S-phase (between the left-side G1 peak and the right-side G2 peak), in contrast to the Halo-tagged Dpb2, which shows an increased proportion of cells in S-phase indicative of slower or problematic DNA replication and an extended S-phase (Fig. S3A). This increased S-phase proportion can also be seen in a *clb5* deletion mutant (YHZ09), which is known to have a lengthier S-phase (Fig. S3B). A small but reproducible change in the pattern, indicative of a longer S-phase, was also observed in the Cdc45-Halo strain (ZEY158). Taken together, the growth curves and flow cytometry data show that the Halo tags and other modifications to our strains do not influence DNA replication and growth and that our measurements and imaging accurately reflect the state of wild-type untagged cells.

**Western blot**

To ensure that the fluorescence tracks seen in imaging were all due to tagged proteins of interest and not to free or cleaved Halo protein, we performed Western blots as described in Supplementary Methods. A different amount was used for each lysate in order to enhance detection of low abundance proteins, prevent saturation by high copy number proteins, and allow easier comparison between both. Since the primary antibody targets an epitope in the Halo tag, we expect to find a band for each protein shifted to a higher molecular weight (due to the added presence of the tag), and to not see any lower molecular weight bands if the Halo tag is not being cleaved. The representative blot (Fig. S4) shows bands of the expected size when including the Halo tag for each of the replisome proteins (Pol12 113kDa, Pri2 96 kDa, Pol2 290kDa, Dpb4 56kDa, Pol3 159kDa, Pol32 75kDa, Mcm4 140kDa, Cdc45 109kDa, Ctf4 139 kDa) and no visible band where free Halo is expected (34kDa), which confirms that the fusions are intact and the Halo tag is properly associated with and not being cleaved from the replisome proteins. The expected molecular weight of HHT1 (Histone H3) with Halo is 50kDa, but other bands can also seen near 30kDa. H3 is known to undergo cleaving at various sites in multiple cellular processes (Yi and Kim, 2018) and we hypothesize the bands we see are not free Halo, but rather these cleaved forms of H3. The absence of any secondary bands in any of the replisome proteins confirms our belief that the Halo tag fusion is stable and the observed fluorescence is indicative of actual replisome dynamics.

**Complications for the use of FRAP to study nuclear events in budding yeast**

There are several limitations and complications associated with performing FRAP and analyzing the results that make it more difficult to interpret, compared to SPT. Some of these issues have previously been reported: nuclear movement, chromosome movement; spatial heterogeneity of binding sites; correct model selection for fitting recovery curves ((Mazza et al., 2012; Mueller et al., 2010)).

Another issue we encountered in our experiments was the small nuclear area relative to the bleaching spot. A fine balance has to be chosen where photobleaching is significant at a particular region, but that does not deplete an important fraction of the total fluorescence in the nucleus –making recovery impossible. This balance is easier to meet in larger nuclei such as that in mammalian cells. However, in yeast, the bleaching area (~1µm diameter) had similar dimensions to the typical nuclei in S-phase (~2µm diameter), making these experiments more complicated. Only a fraction of the experiments was analyzable because of this limitation.

In addition, as noted in the main text, we were unable to determine residence times of the polymerases, likely because of their low chromatin-bound fraction relative to their diffusive fraction. Indeed, in our experiments we were unable to detect the area bleached based on the fluorescence distribution of these subunits (Figure S14), suggesting that the vast majority of the copies of these subunits were diffusive. In contrast, this is not a limitation for sinlge-molecule experiments, since activation of a limited number of copies would help uncover the small fraction of bound molecules.

Figure S1_

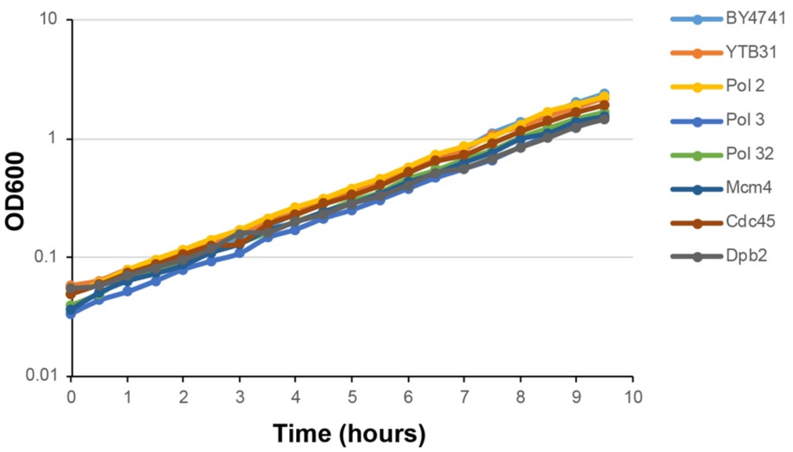

**Generation times in minutes**

| Strain | Repeat 1 | Repeat 2 | Average | StDev |
|---|---|---|---|---|
| BY4741 | 99 | 108 | **104** | 4.4 |
| YTB31 | 104 | 110 | **107** | 2.9 |
| Pol2 | 103 | 103 | **103** | 0.3 |
| Pol3 | 105 | 111 | **108** | 3.2 |
| Pol32 | 105 | 114 | **109** | 4.7 |
| Mcm4 | 107 | 112 | **109** | 2.5 |
| Cdc45 | 106 | 103 | **105** | 1.1 |
| Ctf4 | 107 | 108 | **108** | 0.6 |
| Dpb2 | 118 | 124 | **121** | 3.2 |
| Pol12 CIP- | 101 | 99 | **100** | 1.1 |
| H3 | 127 | 126 | **127** | 0.6 |

**Figure S1. Generation times of Halo-tagged strains used for imaging**. Cultures were diluted from exponential parental cultures and grown in SC at 30°C. OD measurements were taken every 30 minutes and used to plot growth curves. The slopes of the curves were used to calculate the generation time of each strain. The representative graph shown here is from one of two independent experiments. The calculated generation times from both repeats are shown in the table along with their average.

Figure S2_

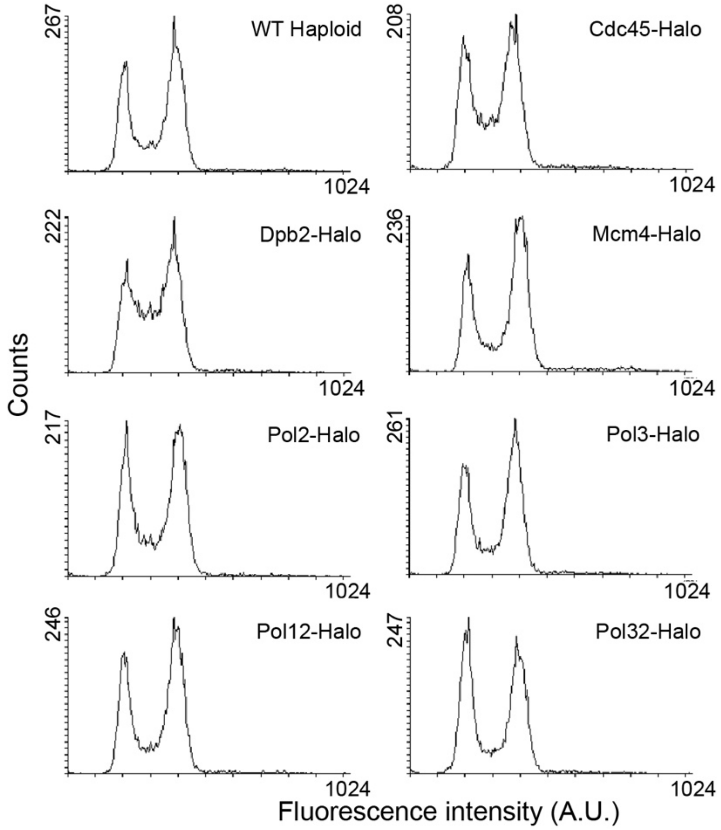

**Figure S2. Flow cytometry analysis of Halo-tagged strains.** Exponential YPD cultures were fixed in 70% ethanol then analyzed by flow cytometry. The cytometer was calibrated using exponentially-growing BY4741 (haploid) and BY4743 (diploid) parental strains. The first peak represents cells with a single chromosomal content (G1), the second peak at double the fluorescence intensity represents cells with fully replicated chromosomes (G2). The area between both peaks represents cell in S-phase. Halo-tagged strains compared to the parental wild-type. All HaloTag fusion have similar profiles to the parental wild-type, except for Dpb2-Halo, which has a higher proportion of cells in S-phase.

Figure S3_

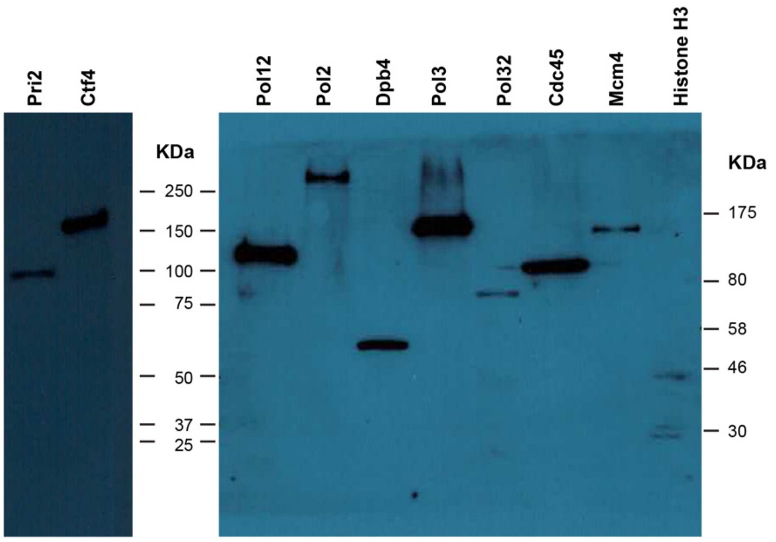

| Protein | Size (kDa) | Size with Halotag (kDa) |
|---|---|---|
| Pol12 | 78 | 113 |
| Pri2 | 62 | 96 |
| Pol2 | 255 | 290 |
| Dpb4 | 22 | 56 |
| Pol3 | 124 | 159 |
| Pol32 | 40 | 75 |
| Mcm4 | 105 | 140 |
| Cdc45 | 74 | 109 |
| Ctf4 | 104 | 139 |
| HHT1 | 15 | 50 |
| Halo alone | | 34 |

**Figure S3. Western blot of HaloTag fusions.** Cell lysates were made from cultures of the various imaging strains. The table shows the size of each protein as well as the expected size when accounting for the presence of the HaloTag. The band for each protein corresponds to the expected size and no smaller bands are seen for the replisome proteins.

Figure S4_

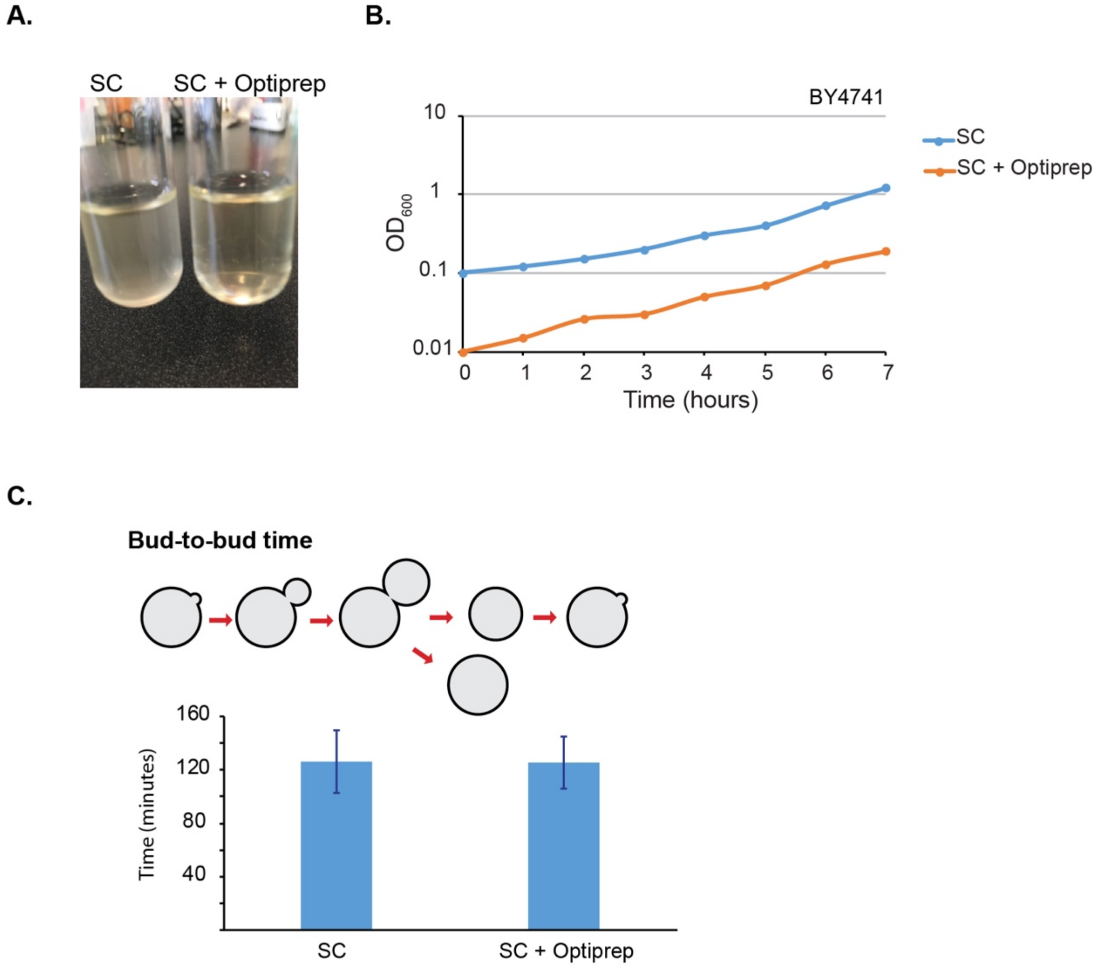

| | SC | SC + Optiprep |
|---|---|---|
| **Mean** | 126.4 | 125.4 |
| **SD** | 23.5 | 19.3 |
| **N** | 31 | 53 |

**Figure S4. Optiprep does not affect cell growth.** A) Addition of Optiprep to an SC culture visibly reduces light absorption by the cells. Both tubes in the image contain two identical halves of the same cultures, with Optiprep added to one of them. The colony counts plated from both cultures are also identical. B) Growth curves of the parental BY4741 strain in SC and SC with 30% Optiprep. Both cultures were diluted from an exponentially growing culture at 30°C and OD measurements were taken every 1h. C) Bud-to-bud measurements on agarose pads. An exponentially growing culture of YTB31 was spotted on SC-agarose pads with the agarose made in either water or Optiprep. The slides were placed on an Olympus microscope at 22°C and bright-field timelapses were taken with 5min intervals for 15h. Single cells were individually timed from the formation of a bud until the formation of the next bud from the cell mother cell and this was used as a measure of doubling time. The mean doubling time for both conditions is shown, with error bars representing standard deviation of the mean.

Figure S5_

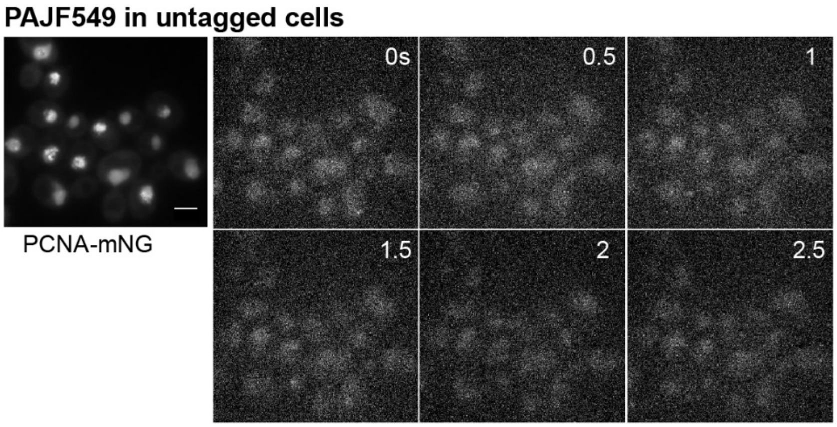

**Figure S5. Background fluorescence in strain carrying no HaloTag fusion.** The strain used was ZEY098, which does not contain a HaloTag fusion. All subsequent analysis – including S-phase nuclei segmentation – was done as previously described.

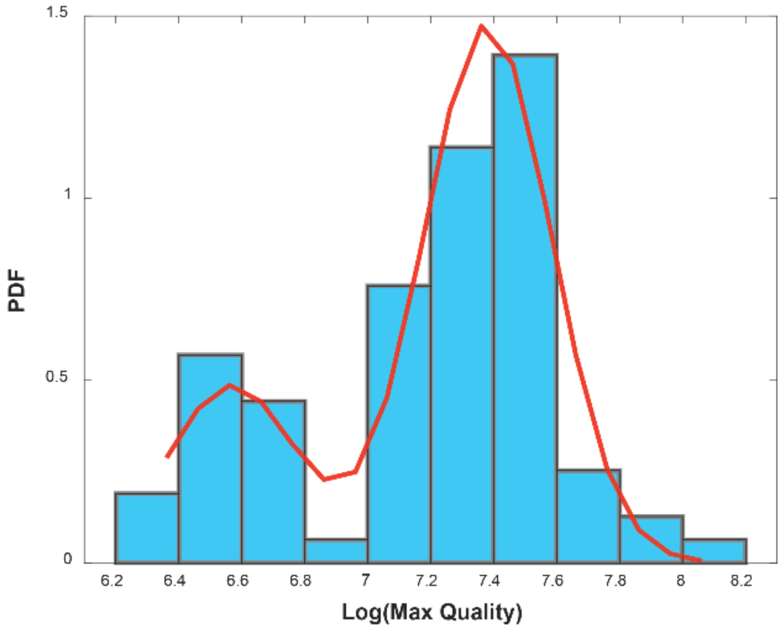

**Figure S6. GMM fitting to quality values**. After the first classification step using Model1, the log-transformed max quality values were fitted with a GMM, and the second largest peak (shown here as being ~ 7.4), was used to scale the quality parameters, prior to classification using Model 2.

Figure S7_

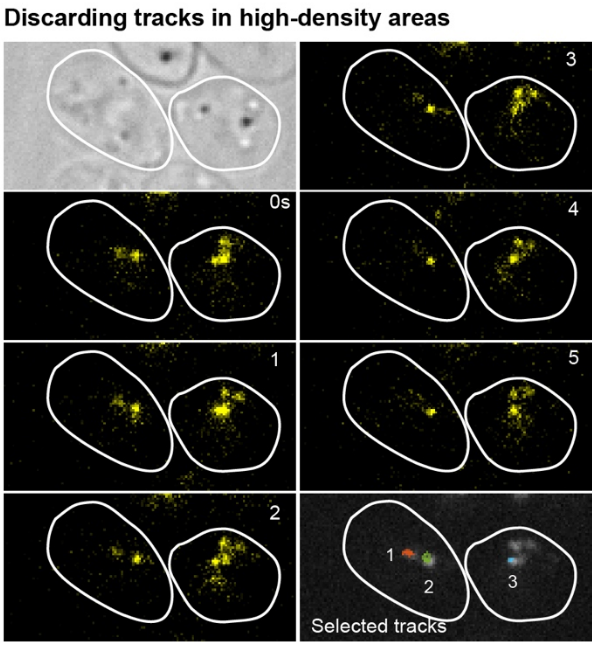

**Figure S7. Discarding tracks in high density regions.** Example of cells showing different densities of activated molecules, taken from Histone H3 data. The cell on the left has a lower density and therefore both tracks (1 and 2) were included in the analysis. In contrast, the cell on the right has a higher density of activated molecules and therefore the majority of tracks were discarded with exception for one track, labelled 3.

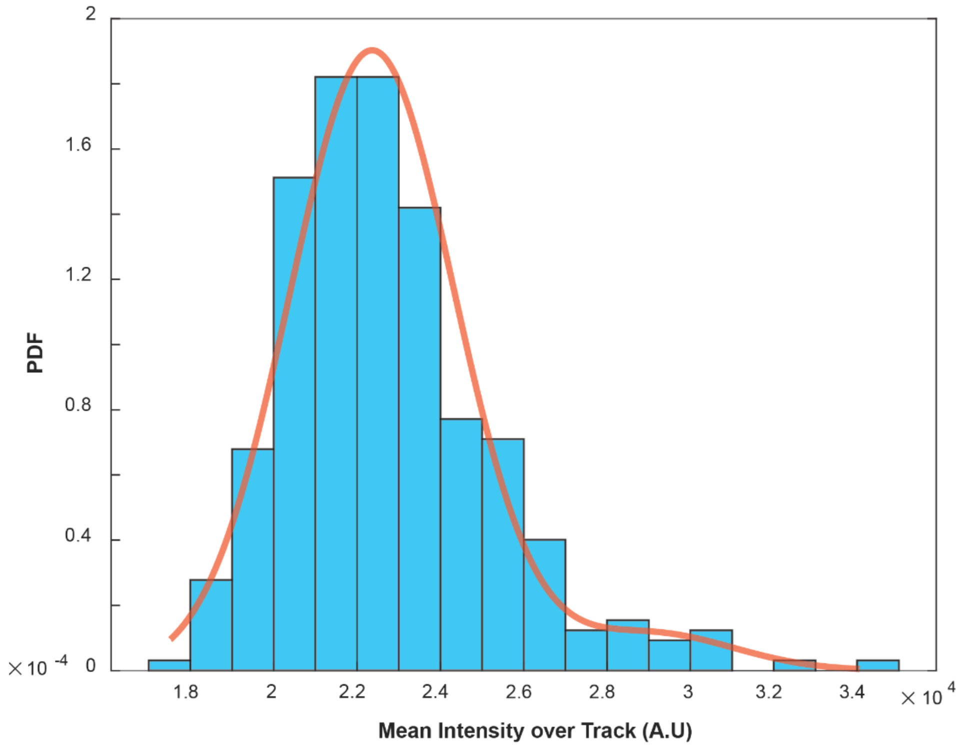

**Figure S8. GMM fitting on mean intensities of tracks.** A GMM was fit to the mean intensities of tracks, with a maximum of 3 components allowed to fit. With the GMM, we clustered the data to isolate tracks representing single molecules.

Figure S9_

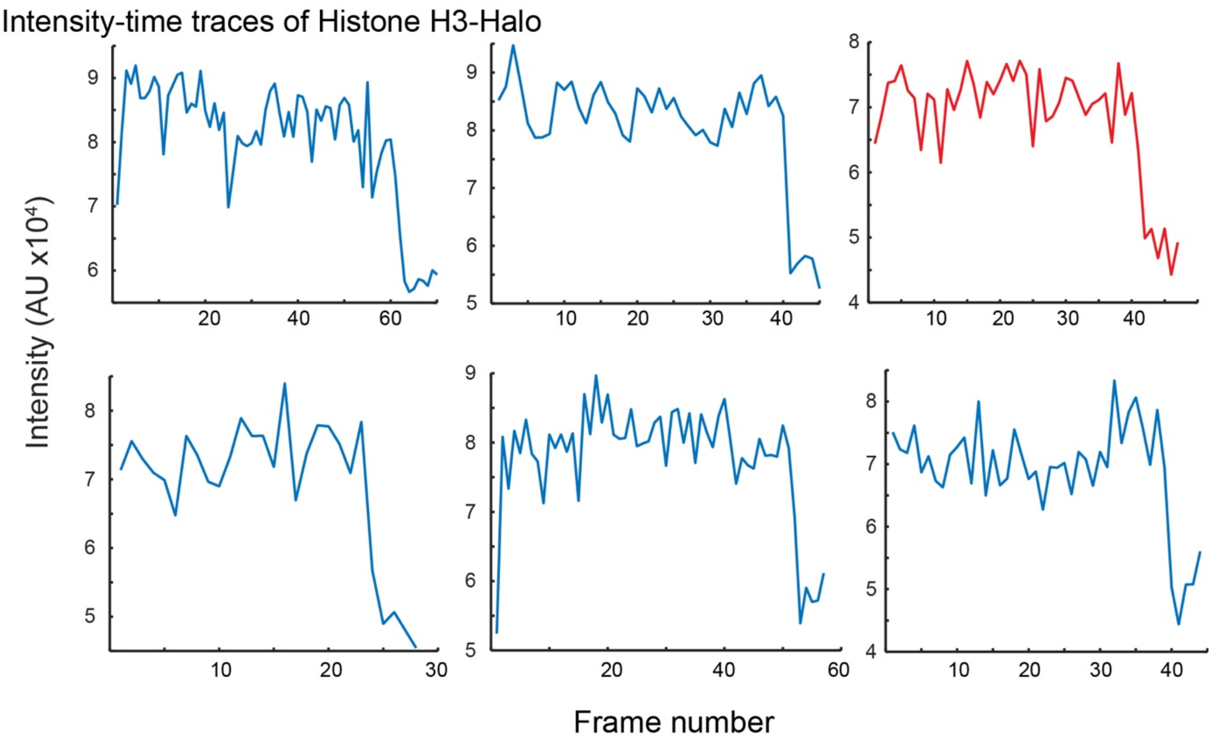

**Figure S9. Examples of single bleaching steps in Histone H3-Halo time traces.** Examples of fluorescence intensity time traces of the H3-Halo show single bleaching steps. Note that they all share similar values in the magnitude of the bleaching step – representing a single molecule equivalent. The trace in orange is the same as in Figure 1G, presented here as comparison.

Figure S10_

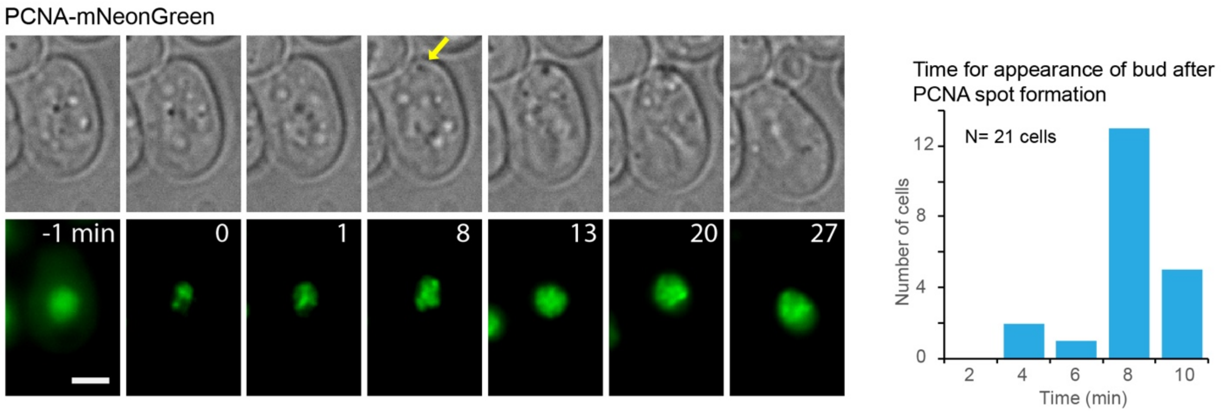

**Figure S10. Correlation of bud and PCNA foci appearance.** An example of the data from 1-minute interval time lapse where it can be observed the appearance of PCNA-mNG spots at 0, followed by the appearance of a bud. The yellow arrow shows the first time point where a noticeable feature, corresponding to the bud, is observed. The plot in the right shows the distribution of times for the appearance of this first feature relative to the appearance of PCNA-mNG spots in multiple cells.

Figure S11_

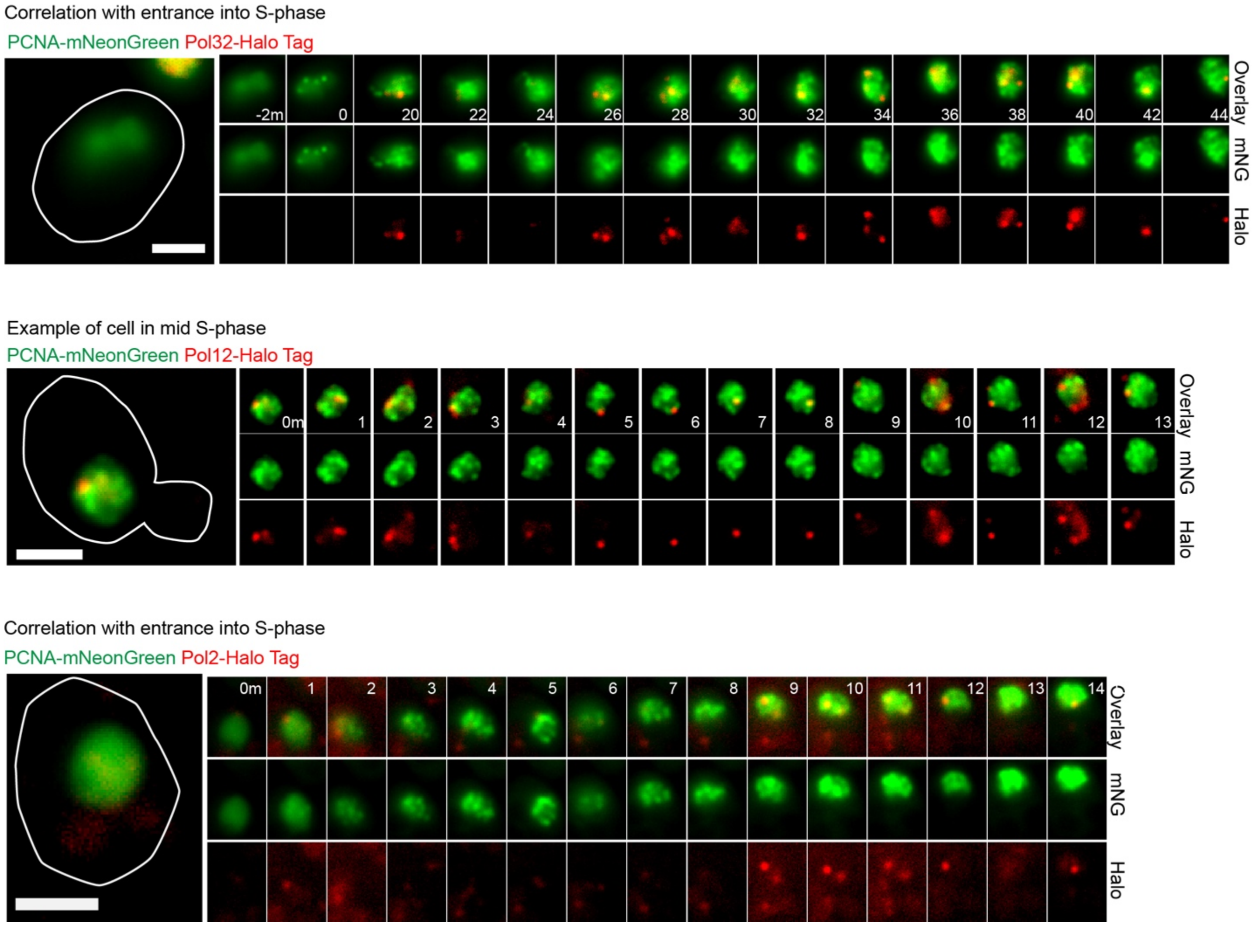

**Figure S11. Examples of correlation between PCNA-mNG and DNA Pol Halo fusions.** Multiple examples showing a correlation between the spatial and temporal distributions of PCNA-mNG and DNA polymerases tagged with HaloTag. Timepoint are in minutes. Scale bars = 2μm.

Figure S12_

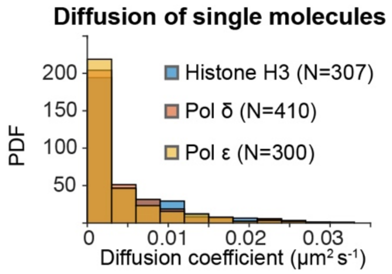

**Figure S12. Distribution of apparent diffusion coefficients of individual tracks.** Comparison of the distributions of apparent diffusion coefficients for 3 different proteins, after estimating the apparent diffusion coefficient from individual tracks classified as being bound with the ML classification.

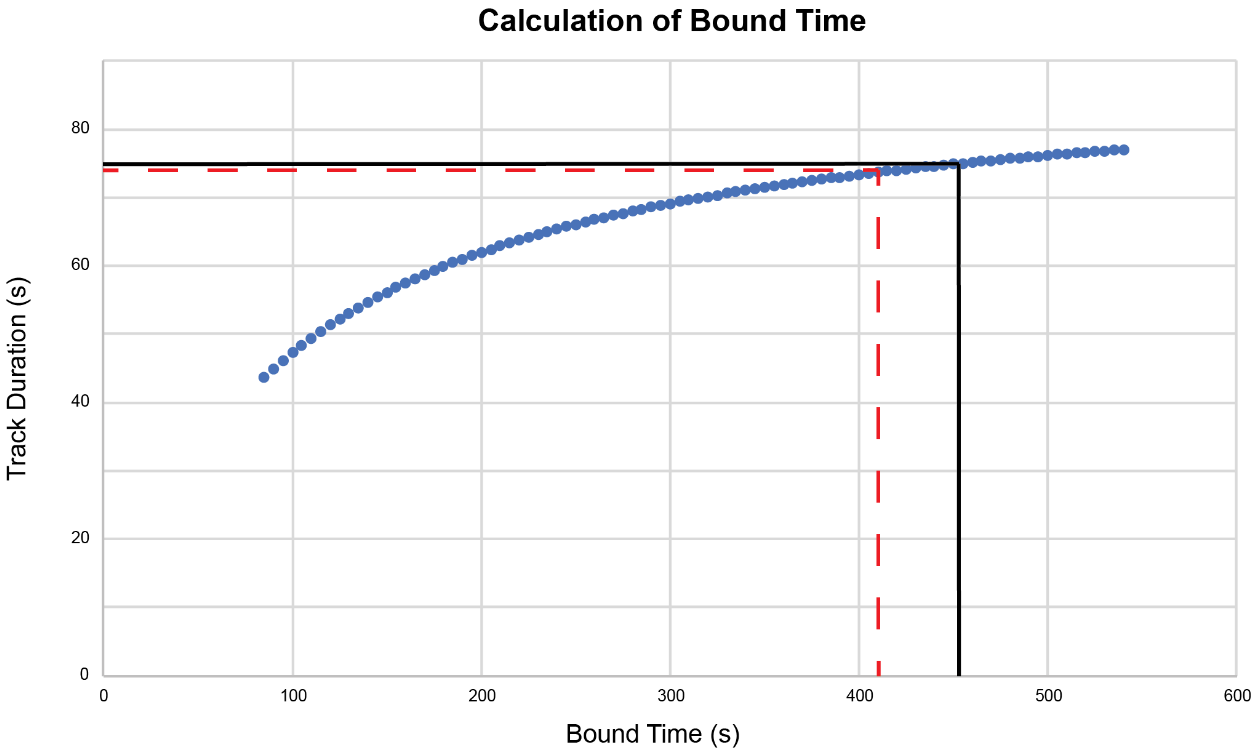

**Figure S13. Calculation of minimum bound time using 8 interval data**. Black solid indicated the bound time value that results in a track duration time equal to the lower bound of the CI for the bleaching data. The red dotted line represents the track duration that is 1s lower than the lower bound of the bleaching estimate CI.

Figure S14_

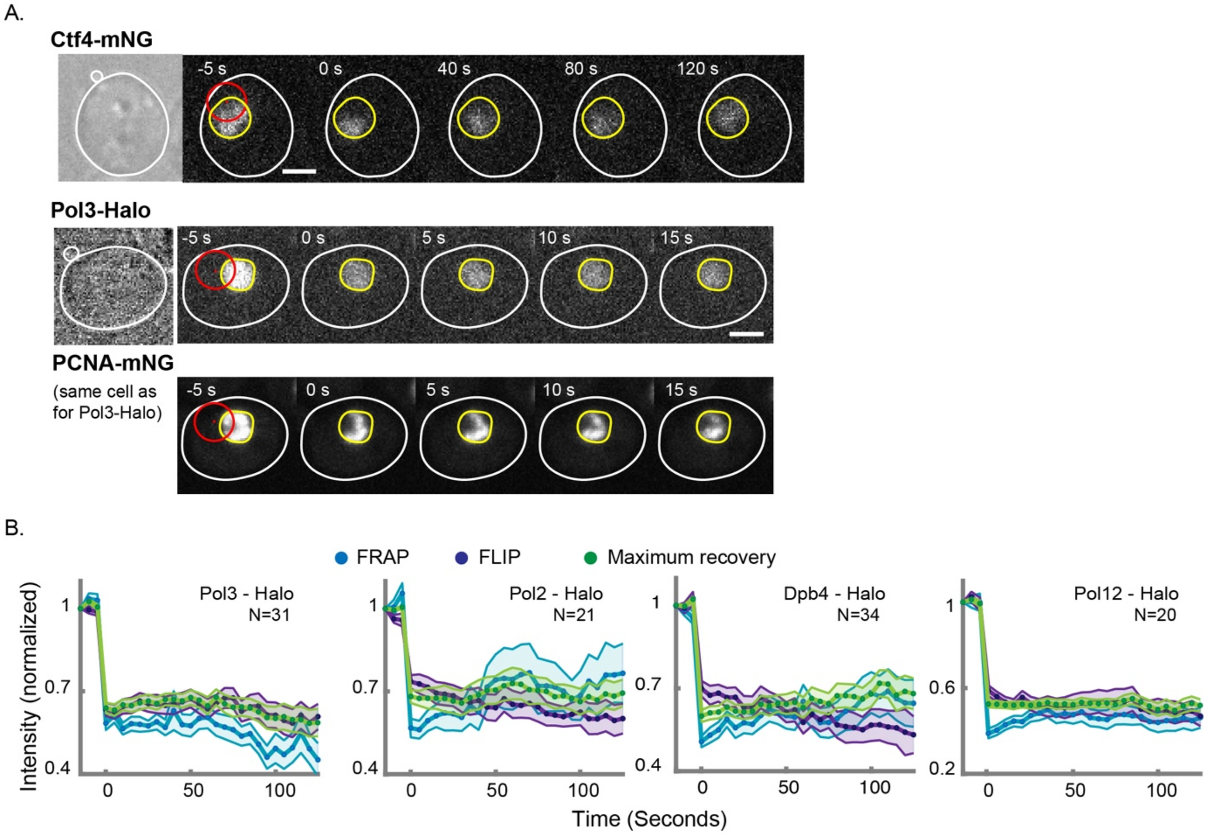

**Figure S14. Examples of FRAP for DNA polymerases.** (**A**) Examples of FRAP experiments for CTF4-mNG and Pol3-Halo. Note that no noticeable bleached area can be observed in the case of Pol3-Halo, but the bleached area is clearly visible in the distribution of PCNA-mNG for the same cell. (B) Plots showing the average intensities over time for different DNA polymerase subunits. Note the similar values of FRAP, FLIP and maximum recovery in the time point immediately after the photobleaching step, suggesting that a very fast redistribution of the fluorescence. Among the polymerases, the results for Pol12-Halo were the most promising. However, we were unable to obtain sensible numbers for it, and is an area for future development.

Figure S15_

**A.**

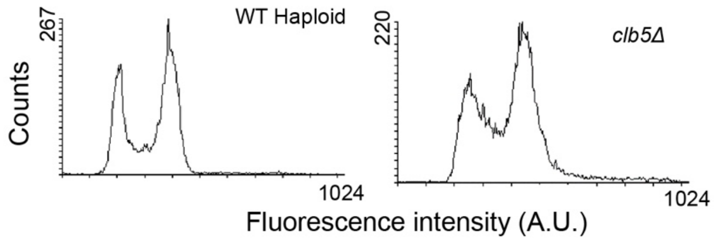

**B.**

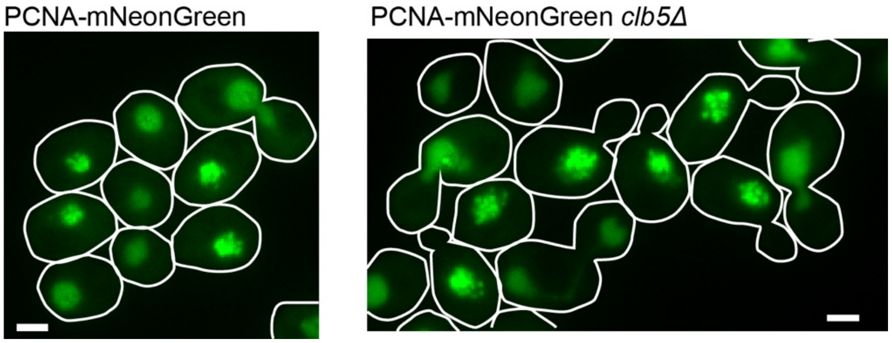

**Figure S15. Characterization of *clb5Δ* PCNA-mNG strain.** (**A**) Flow cytometry for the wt BY4741 haploid strain and for a strain carrying Pol32-Halo PCNA-mNG *clb5Δ*. Note the higher values between the 1C and 2C peaks, representing a greater fraction of cells undergoing DNA replication. (**B**) Distribution of fluorescence in cells carrying PCNA-mNG and either a wt allele for *clb5* or a *clb5Δ*. Note the higher frequency of cells with PCNA spots and the apparent lower density of spots in the *clb5Δ* mutant. The locations of spots also seemed more dispersed in the *clb5Δ* mutant.

| Strain | Genotype |
|---|---|
| **BY4741** | MATa his3Δ1 leu2Δ0 met15Δ0 ura3Δ0 |
| **BY4742** | MATα his3Δ1 leu2Δ0 lys2Δ0 ura3Δ0 |
| **BY4743** | MATa/α his3Δ1/his3Δ1 leu2Δ0/leu2Δ0 LYS2/lys2Δ0 met15Δ0/MET15 ura3Δ0/ura3Δ0 |
| **YHZ09** | MATa his3Δ1 leu2Δ0 LYS2 met15Δ0 ura3Δ0 clb5Δ0::KanMX POL30-mNeonGreen-Nat |
| **YTB31** | MATα his3Δ1 leu2Δ0 lys2Δ0 ura3Δ0 POL30-mNeonGreen-Nat |
| **YTK1414** | MATa/α his3Δ1/his3Δ1 leu2Δ0/leu2Δ0 LYS2/lys2Δ0 met15Δ0/MET15 ura3Δ0/ura3Δ0 PDR5/pdr5Δ::KanMX |
| **ZEY098** | MATa his3Δ1 leu2Δ0 LYS2 met15Δ0 ura3Δ0 pdr5Δ0::KanMX POL30-mNeonGreen-Nat |
| **YJL10** | MATα his3Δ1 leu2Δ0 lys2Δ0 MET15 ura3Δ0 pdr5Δ0::KanMX POL30-mNeonGreen-Nat POL12-Halo-HygB |
| **ZEY136** | MATa his3Δ1 leu2Δ0 LYS2 met15Δ0 ura3Δ0 pdr5Δ0::KanMX POL30-mNeonGreen-Nat PRI2-Halo-HygB |
| **YJL02** | MATα his3Δ1 leu2Δ0 lys2Δ0 MET15 ura3Δ0 pdr5Δ0::KanMX POL30-mNeonGreen-Nat POL2-Halo-HygB |
| **YJL18** | MATα his3Δ1 leu2Δ0 lys2Δ0 MET15 ura3Δ0 pdr5Δ0::KanMX POL30-mNeonGreen-Nat DPB4-Halo-HygB |
| **YJL24** | MATα his3Δ1 leu2Δ0 lys2Δ0 MET15 ura3Δ0 pdr5Δ0::KanMX POL30-mNeonGreen-Nat POL3-Halo-HygB |
| **ZEY057** | MATα his3Δ1 leu2Δ0 lys2Δ0 MET15 ura3Δ0 pdr5Δ0::KanMX POL30-mNeonGreen-Nat POL3-Halo-HygB clb5Δ0::KanMX |
| **YJL11** | MATα his3Δ1 leu2Δ0 lys2Δ0 MET15 ura3Δ0 pdr5Δ0::KanMX POL30-mNeonGreen-Nat POL32-Halo-HygB |
| **YAY256** | MATa his3D1 leu2D0 lys2D0 MET15 ura3D0 pdr5Δ::KanMX POL30-mNeonGreen-Nat MCM4-Halotag-HygB |
| **ZEY158** | MATα his3Δ1 leu2Δ0 lys2Δ0 MET15 ura3Δ0 pdr5Δ0::KanMX POL30-mNeonGreen-Nat CDC45-Halo-HygB |
| **ZEY077** | MATα his3Δ1 leu2Δ0 lys2Δ0 MET15 ura3Δ0 pdr5Δ0::KanMX POL30-mNeonGreen-Nat CTF4-Halo-HygB |
| **ZEY206** | MATa his3Δ1 leu2Δ0 LYS2 met15Δ0 ura3Δ0 pdr5Δ0::KanMX POL30-mNeonGreen-Nat RFA1-Halo-HygB |
| **YAJ05** | MATα his3Δ1 leu2Δ0 lys2Δ0 MET15 ura3Δ0 pdr5Δ0::KanMX POL30-mNeonGreen-Nat POL12-Halo-HygB Pol14A-URA3 |
| **YTK1434** | MATa his3Δ1 leu2Δ0 met15Δ0 ura3Δ0 pdr5Δ0::KanMX HHT1-Halo-URA3 |
| **YTB56** | MATa/α his3Δ1/his3Δ1 leu2Δ0/leu2Δ0 LYS2/lys2Δ0 met15Δ0/MET15 ura3Δ0/ura3Δ0 MCM4-mNeonGreen-Nat/MCM4-mNeonGreen-Nat |
| **YTB57** | MATa/α his3Δ1/his3Δ1 leu2Δ0/leu2Δ0 LYS2/lys2Δ0 met15Δ0/MET15 ura3Δ0/ura3Δ0 CTF4-mNeonGreen-Nat/CTF4-mNeonGreen-Nat |
| **ZEY202** | MATa/α his3Δ1/his3Δ1 leu2Δ0/leu2Δ0 LYS2/lys2Δ0 met15Δ0/MET15 ura3Δ0/ura3Δ0 RFA1-mNeonGreen-Nat/RFA1-mNeonGreen-Nat |

**Table S1. Strains used.** The *S. cerevisiae* strains used in this study are listed along with their genotypes.

| Primer | Description | Sequence |
|---|---|---|
| **TB81** | C-terminal mNeonGreen tagging of PCNA (F) | cctacagtttttcttggctcctaaatttaatgacgaagaaGGTGACGGTGCTGGTTTA |
| **TB82** | C-terminal mNeonGreen tagging of PCNA (R) | tttattatttttagtatacaactatatagataatttacatCACAGGAAACAGCTATGACC |
| **TB98** | Screen C-terminal tag of PCNA (F) | AGAGTTGGTATCAGGCTCTC |
| **TB99** | Screen C-terminal tag of PCNA (R) | AAGCTGATATTTAACGCATCTTAG |
| **TB123** | C-terminal Halo tagging of Pol12 (F) | caacgtgtggaagcgcgctagagttgacttgattgctagtGGTGACGGTGCTGGTTTA |
| **TB124** | C-terminal Halo tagging of Pol12 (R) | accttgagctattccattagttaagtttgaattaaatataCACAGGAAACAGCTATGACC |
| **AY9** | Screen C-terminal tag of Pol12 (F) | CTTGTTGAAGGTGAAGAGCC |
| **AY10** | Screen C-terminal tag of Pol12 (R) | GCCAGTTTCAAGGTCGATAG |
| **TB125** | C-terminal Halo tagging of Pri2 (F) | gaagctggaaaaggaaaaactattcaataatggtaatcatGGTGACGGTGCTGGTTTA |
| **TB126** | C-terminal Halo tagging of Pri2 (R) | tttagttatctcttcgctttttttccttttccctttctgcaCACAGGAAACAGCTATGACC |
| **AY11** | Screen C-terminal tag of Pri2 (F) | CGAAAGATCAAGGCAACTGC |
| **AY12** | Screen C-terminal tag of Pri2 (R) | TTTTTGACCATACTTACAGTAGAC |
| **TB61** | C-terminal Halo tagging of Pol2 (F) | ttttgatatattattgagttgtattgctgatttgaccataGGTGACGGTGCTGGTTTA |
| **TB62** | C-terminal Halo tagging of Pol2 (R) | ggtaaagaggccattgaacctcgcgttatatactgcttacCACAGGAAACAGCTATGACC |
| **TB44** | Screen C-terminal tag of Pol2 (F) | TGCCCACTGTCCATGTGC |
| **TB45** | Screen C-terminal tag of Pol2 (R) | CAACTTCCGGAGTGGTCAC |
| **AY23** | C-terminal Halo tagging of Dbp4 (F) | ccaagatgtagaaactagagttcaaaaccttgagcaaacgGGTGACGGTGCTGGTTTA |
| **AY24** | C-terminal Halo tagging of Dbp4 (R) | gagtggtggcaagcactactagacagtttccatagcggggCACAGGAAACAGCTATGACC |
| **AY43** | Screen C-terminal tag of Dpb4 (F) | AAGGCGATGCATTACAGGAC |
| **AY44** | Screen C-terminal tag of Dpb4 (R) | TTCCCCGGCTTGCAAATAAC |
| **TB63** | C-terminal Halo tagging of Pol3 (F) | aaaagagctgcaggagaaagtagaacaattaagcaaatggGGTGACGGTGCTGGTTTA |
| **TB64** | C-terminal Halo tagging of Pol3 (R) | cctttcttaatcctaatatgatgtgccaccctatcgttttCACAGGAAACAGCTATGACC |
| **TB71** | Screen C-terminal tag of Pol3 (F) | GCGCTGGTAACTTACATAGTG |
| **TB72** | Screen C-terminal tag of Pol3 (R) | TGAATCTGGATTTTTCCAAGTATC |
| **AY27** | C-terminal Halo tagging of Pol32 (F) | gcaaggaacattggaaagctttttcaaaagaaaggcaaaaGGTGACGGTGCTGGTTTA |
| **AY28** | C-terminal Halo | tcacaattagtaatggaaagtgtttggaaaaaaaagaagaCACAGGAAACAGCTATGACC |

| | | |
|---|---|---|
| | tagging of Pol32 (R) | |
| **AY47** | Screen C-terminal tag of Pol32 (F) | AAGCAAGAAACGCCGTCATC |
| **AY48** | Screen C-terminal tag of Pol32 (R) | TTCTATCACGTAAGTTGACATTTG |
| **AY29** | C-terminal Halo/PCNA tagging of Mcm4 (F) | cgagggtgtaaggagatcagttcgcctgaataaccgtgtcGGTGACGGTGCTGGTTTA |
| **AY30** | C-terminal Halo/PCNA tagging of Mcm4 (R) | ttattaattgttacgcagggaatgattgtagtagacagcaCACAGGAAACAGCTATGACC |
| **AY49** | Screen C-terminal tag of Mcm4 (F) | GGAAGCCTTGTCAAGATTGC |
| **AY50** | Screen C-terminal tag of Mcm4 (R) | ATCGAGCCTACATACAGTATTG |
| **TB87** | C-terminal Halo tagging of Cdc45 (F) | ttcaccattcctggagaagctgaccttgagtggattgttaGGTGACGGTGCTGGTTTA |
| **TB88** | C-terminal Halo tagging of Cdc45 (R) | tatgctggtatatatgtacgactaaataatataaatttgaCACAGGAAACAGCTATGACC |
| **TB104** | Screen C-terminal tag of Cdc45 (F) | AAATAACTGCAGAAACGGATGC |
| **TB105** | Screen C-terminal tag of Cdc45 (R) | AGAGCCGCGCACAAAATATG |
| **TB59** | C-terminal Halo/PCNA tagging of Ctf4 (F) | taataatataagggaagctagatatgaacagcaattgaaaGGTGACGGTGCTGGTTTA |
| **TB60** | C-terminal Halo/PCNA tagging of Ctf4 (R) | tcaaataattgtctcttgcgtatatatattttacatttttCACAGGAAACAGCTATGACC |
| **TB69** | Screen C-terminal tag of Ctf4 (F) | CACTTACTGCAGCCGTTAAG |
| **TB70** | Screen C-terminal tag of Ctf4 (R) | TAATGTGGGAGCATTTTGAACG |
| **TB115** | C-terminal Halo/PCNA tagging of Rfa1 (F) | cgactatcttgccgatgagttatccaaggctttgttagctGGTGACGGTGCTGGTTTA |
| **TB116** | C-terminal Halo/PCNA tagging of Rfa1 (R) | tatgttacatagattaaatagtacttgattatttgatacaCACAGGAAACAGCTATGACC |
| **AY01** | Screen C-terminal tag of Rfa1 (F) | CAGCTTGAATTACAGGGCTG |
| **AY02** | Screen C-terminal tag of Rfa1 (F) | CCGCCCTTCAAAAACTTGAC |
| **NK57** | Sequence mutations in POL1-4A CIP mutant (F) | ATATACGACGAAATCGACG |
| **NK58** | Sequence mutations in POL1-4A CIP mutant (R) | CCACATCATCCAATAAATCC |

**Table S2.** Primers used in this study, with a short description and their sequence.

| Protein | Time Interval (s) | N | Tbound Alpha estimate | Tbound Beta estimate | BIC Conclusion | LLR Test P value | Estimates as bounds | Chi2GoF P values |
|---|---|---|---|---|---|---|---|---|
| Pol32 | 0.5 | 163 | 899.91 | 0.1 | Two-exponential | 3.31*10^-6 | Yes | 0.31 |
| Pol32 | 1 | 415 | 899.86 | 61.39 | Single-Exponential | 0.67 | Yes | 0.22 |
| Pol3 | 1 | 324 | 899.99 | 34.64 | Single-Exponential | 0.17 | Yes | 0.29 |
| Pol12 | 1 | 253 | 255.9 | 16.48 | Single-Exponential | 0.17 | No | 0.0689 |
| Pri2 | 1 | 118 | 31.7 | 899.9364 | Single-Exponential | 0.7139 | Yes | 0.0027 |
| Pol12 | 8 | 95 | 87.91 | 30.28 | Single-Exponential | 1 | Yes | 0.1821 |

**Table S3**. Results from statistical tests to test for two-exponential behavior on combined data sets. The lower and upper bounds on the estimates were 0.1s and 900s, respectively. The bleach time was 15s, 23s, and 90, for the 0.5s, 1s, and 8s time interval, respectively. We used the BIC test and LLR test, under the null hypothesis of single-exponential distribution, to compare whether a two-exponential (more complex) model significantly fits better than the single-exponential. If the estimates were simply the bounds, it indicated that the algorithm could not find two different behaviours within the bleaching time. We also performed the chi square goodness of fit test, under the null hypothesis that the distribution comes from a single-exponential distribution.

| Replisomes | Excess Copy Number | D_Pol Delta (um^2/s) | D_Replisome (um^2/s) | Initial Bound Fraction | Ttrack (s) | Single or Two Exponential |
|---|---|---|---|---|---|---|
| 300 | 1600 | 0.5 | 0.005 | 0.25 | 12.89[8.90,16.88] | Single |
| 300 | 1600 | 0.5 | 0.005 | 0.75 | 11.82[8.16,15.48] | Single |
| 300 | 1600 | 0.5 | 0.05 | 0.25 | 14.52[10.02,19.01] | Single |
| 300 | 1600 | 0.5 | 0.05 | 0.75 | 12.47[8.60, 16.33] | Single |
| 300 | 1600 | 5 | 0.005 | 0.25 | 12.28[8.48,16.09] | Single |
| 300 | 1600 | 5 | 0.005 | 0.75 | 11.80[8.14,15.45] | Single |
| 300 | 1600 | 5 | 0.05 | 0.25 | 11.07[7.64,14.51] | Single |
| 300 | 1600 | 5 | 0.05 | 0.75 | 11.35[7.83,14.86] | Single |

**Table S4.** Results of simulations testing for rebinding through a range of parameters, along with outcome from statistical tests to check if two-exponential model fits better than single-exponential.

| Replisomes | Excess Copy Number | D_Pol Delta (um^2/s) | D_Replisome (um^2/s) | Mean Fraction of Time Bound by Pol Delta |
|---|---|---|---|---|
| 300 | 1600 | 0.5 | 0.005 | 0.2909 |
| 300 | 1600 | 0.5 | 0.05 | 0.2982 |
| 300 | 1600 | 5 | 0.005 | 0.2977 |
| 300 | 1600 | 5 | 0.05 | 0.297 |
| 300 | 3200 | 0.5 | 0.005 | 0.3526 |
| 300 | 3200 | 0.5 | 0.05 | 0.3406 |
| 300 | 3200 | 5 | 0.005 | 0.3487 |
| 300 | 3200 | 5 | 0.05 | 0.3523 |

**Table S5 .** Results of simulations testing for replisome occupancy through a range of parameters.

| | Mean Track Duration (s) | Std.Error (s) | CI (s) | N |
|---|---|---|---|---|
| **0.5s** | | | | |
| Pol32 | 13.23 | 1.11 | [11.3,15.59] | 163 |
| Rfa1 | 5.06 | 0.38 | [4.45,5.96] | 184 |
| Rfa1 with HU | 7.22 | 0.52 | [6.29, 8.38] | 224 |
| **1.0s** | | | | |
| Cdc45 | 23.24 | 1.2 | [21.10,25.75] | 406 |
| Dpb4 | 20.84 | 1.38 | [18.28, 23.60] | 269 |
| Pol3 | 22.32 | 1.33 | [19.76, 24.94] | 324 |
| Pol32 | 24.51 | 1.26 | [ 22.25, 27.10] | 415 |
| Pol12 | 17.09 | 1.12 | [15.05, 19.30] | 253 |
| Pol2 | 20.11 | 1.01 | [18.31, 22.27] | 305 |
| Ctf4 | 24.23 | 1.08 | [22.36,26.63] | 384 |
| Pri2 | 14.75 | 1.33 | [12.58, 18.00] | 118 |
| Po12-CIP- | 15.88 | 1.09 | [14.02, 18.18] | 240 |
| ΔClb5-Pol3 | 23.75 | 2 | [20.25,28.22] | 114 |
| **8s Interval** | | | | |
| Cdc45 | 84.16 | 7.71 | [70.29, 101.73] | 129 |
| Dpb4 | 66.94 | 6.06 | [56.64, 80.89] | 128 |
| Pol32 | 65.54 | 5.61 | [56.06, 78.09] | 130 |
| Pol12 | 43.96 | 4.64 | [36.49, 55.61] | 95 |
| Ctf4 | 69.61 | 5.18 | [60.09, 80.50] | 147 |
| Histone H3 | 89.52 | 5.33 | [80.01, 100.44] | 263 |
| Pol12-CIP- | 25.41 | 3.89 | [18.73,34.68] | 85 |
| **20s Interval** | | | | |
| Cdc45 | 178.44 | 18.24 | [143.61, 217.19] | 64 |
| Dpb4 | 128.91 | 11.95 | [107.83, 155.94] | 92 |
| Pol3 | 104.55 | 13.34 | [81.36,136.01] | 44 |
| Pol32 | 102.27 | 13.89 | [77.73, 135.32] | 44 |
| Histone H3 | 153.67 | 15.29 | [126.33, 185.26] | 60 |

**Table S6. Track durations from combined data sets.** Track durations from different experiments collected with the same time interval were combined into a single data set, which was then fitted with a truncated exponential model. Standard errors and CI were calculated through bootstrapping 1000 samples.

| | Bound Time Estimate (s) | Std error (s) | CI (s) |
|---|---|---|---|
| **0.5s** | | | |
| Rfa1 | 8.63 | 1.10 | [6.66,11.03] |
| Rfa1 + HU | 17.62 | 2.88 | [13.05,24.32] |
| **1s** | | | |
| Pol12 | 65.53 | 15.29 | [ 45.971, 107.24] |
| Pri2 | 40.74 | 9.97 | [27.38, 71.64] |
| Pol12-CIP | 50.71 | 11.32 | [34.34, 73.10] |
| **8s** | | | |
| Dpb4 | 265.39 | 141.02 | [138.89, 625.63] |
| Pol32 | 244.67 | 88.30 | [148.90, 486.98] |
| Pol12 | 86.38 | 17.01 | [60.60,127.43] |
| Ctf4 | 312.98 | 164.77 | [186.9, 1037.3] |
| Pol12-CIP- | 35.48 | 7.93 | [23.57,57.55] |
| **20s** | | | |
| Dpb4 | 800.06 | 584.62 | [405.30, 2531.00] |
| Pol32 | 305.76 | 124.65 | [155.99, 660.27] |
| Pol3 | 327.08 | 128.16 | [170.05, 674.19] |

**Table S7. Bound times for different subunits.** Bound time estimates from combined data sets shown in Table S6, for cases where the track duration estimates were different than the photobleaching time estimates.. Errors were calculated using 1000 bootstrap samples.

| Date (year/month/day) | Protein | Sample Size | Mean Track Duration (s) [95% CI] |
|---|---|---|---|
| **500ms Interval** | | | |
| 20181206 | Pol32 | 119 | 12.91 [10.59, 15.23] |
| 20181122 | Pol32 | 44 | 14.11 [9.94, 18.28] |
| 20180511 | Histone H3 | 126 | 15.69[12.95, 18.44] |
| 20200129 | Histone H3 | 244 | 12.23s [10.7, 13.77] |
| 20200115 | Rfa1 | 54 | 6.1 [4.47, 7.73] |
| 20200116 | Rfa1 | 114 | 5.00 [4.08, 5.97] |
| 20200203 | Rfa1 | 70 | 5.16 [3.95, 6.37] |
| 20200204 | Rfa1 + HU | 47 | 6.43 [4.59, 8.26] |
| 20200206 | Rfa1 + HU | 68 | 6.65 [5.07, 8.23] |
| 20200207 | Rfa1 + HU | 109 | 7.91 [6.42, 9.39] |
| **1s Interval** | | | |
| 20180417 | Cdc45 | 160 | 23.55[19.90, 27.2] |
| 20180427 | Cdc45 | 173 | 22.06[18.77, 25.34] |
| 20180830 | Cdc45 | 73 | 25.37[19.55, 31.19] |
| 20180810 | Mcm4 | 95 | 27.76[22.18, 33.34] |
| 20180412 | Pol2 | 107 | 20.73 [16.8, 24.66] |
| 20180802 | Pol2 | 120 | 18.75[15.4, 22.1] |
| 20180803 | Pol2 | 78 | 21.37[16.63, 26.11] |
| 20180808 | Dpb4 | 88 | 20.16[15.95, 24.37] |
| 20180424 | Dpb4 | 181 | 21.17[18.08, 24.25] |
| 20180621 | Pol3 | 138 | 22.8 [19, 26.61] |
| 20180622 | Pol3 | 39 | 33.15 [22.75, 43.56] |
| 20180724 | Pol3 | 69 | 15.84 [12.1, 19.58] |
| 20190429 | Pol3 | 78 | 21.8 [16.95, 26.63] |
| 20180406 | Pol32 | 182 | 26.95[23.04, 30.87] |
| 20180429 | Pol32 | 127 | 23.17 [19.14, 27.2] |
| 20180809 | Pol32 | 106 | 21.94 [17.77, 26.12] |
| 20190531 | ΔClb5-Pol3-Halo | 52 | 25.06 [18.25, 31.87] |
| 20190602 | ΔClb5-Pol3-Halo | 62 | 22.65 [17.0, 28.28] |
| 20180618 | Pol12 | 54 | 19.81[14.53, 25.1] |
| 20180626 | Pol12 | 140 | 16.67 [13.91, 19.43] |
| 20190108 | Pol12 | 59 | 15.59[11.61, 19.57] |
| 20190713 | Pri2 | 45 | 17.31 [12.25, 22.37] |
| 20190718 | Pri2 | 73 | 13.18 [10.16, 16.2] |
| 20190806 | Pol12-CIP- | 56 | 18.96 [14, 23.92] |
| 20190807 | Pol12-CIP- | 77 | 15.84 [12.31, 19.39] |
| 20190813 | Pol12-CIP- | 49 | 15.26 [10.99, 19.54] |
| 20190814 | Pol12-CIP- | 58 | 13.48 [10.01. 16.95] |
| 20190111 | Ctf4 | 306 | 23.72 [21.06, 26.38] |
| 20190112 | Ctf4 | 78 | 26.23 [20.41, 32.05] |
| 20180904 | Histone H3 | 471 | 23.12[21.03, 25.21] |
| 20190627 | Histone H3 | 113 | 23.09 [18.84, 27.36] |
| **8s Interval** | | | |
| 20180906 | Cdc45 | 16 | 76.5 [39.02, 113.98] |
| 20180913 | Cdc45 | 14 | 58.86 [28.03, 89.69] |
| 20180927 | Cdc45 | 13 | 53.54 [24.44, 82.64] |
| 20181005 | Cdc45 | 68 | 92.23 [70.32, 114.16] |
| 20181125 | Cdc45 | 18 | 102.22 [55, 149.45] |
| 20180817 | Dpb4 | 53 | 53.28 [38.94, 67.63] |
| 20180921 | Dpb4 | 44 | 58.36 [41.12, 75.61] |
| 20181204 | Dpb4 | 31 | 102.45 [66.39, 138.52] |
| 20180827 | Pol32 | 31 | 72.26 [46.82, 97.69] |
| 20180919 | Pol32 | 22 | 53.45 [31.12, 75.79] |
| 20181130 | Pol32 | 36 | 75.11 [50.57, 99.65] |
| 20190216 | Pol32 | 41 | 58.54 [40.62, 76.45] |
| 20180914 | Pol12 | 12 | 70 [30.39, 109.61] |
| 20180925 | Pol12 | 28 | 28.29 [17.81, 38.76] |
| 20181129 | Pol12 | 15 | 51.2 [25.29, 77.11] |
| 20190109 | Pol12 | 12 | 34.67 [15.05, 54.28] |
| 20190129 | Pol12 | 28 | 48.57 [30.58, 66.56] |
| 20200212 | Pol12-CIP- | 16 | 29.00 [14.79, 43.21] |
| 20200222 | Pol12-CIP- | 12 | 45.33 [19.69, 70.98] |
| 20200226 | Pol12-CIP- | 19 | 26.95 [14.83, 39.06] |
| 20200305 | Pol12-CIP- | 20 | 15.6 [8.76, 22.44] |
| 20200310 | Pol12-CIP- | 18 | 18.22 [10.22, 31.88] |
| 20190115 | Ctf4 | 73 | 68.6 [52.86, 84.34] |
| 20190116 | Ctf4 | 74 | 70.59 [54.51, 86.68] |
| 20180822 | Histone H3 | 155 | 79.33 [66.84, 91.82] |
| 20180907 | Histone H3 | 108 | 104.15 [84.51, 123.79] |
| **20s Interval** | | | |
| 20181009 | Cdc45 | 15 | 192 [94.84, 289.16] |
| 20181030 | Cdc45 | 18 | 147.78 [79.51, 216.05] |
| 20181107 | Cdc45 | 31 | 189.68 [122.91, 256.45] |
| 20181017 | Dpb4 | 21 | 153.33 [87.75, 218.91] |
| 20181102 | Dpb4 | 51 | 114.12 [82.8, 145.44] |
| 20181113 | Dpb4 | 20 | 141 [79.21, 202.79] |
| 20181011 | Pol32 | 16 | 87 [44.63, 130.37] |
| 20181031 | Pol32 | 15 | 128 [63.22, 192.78] |
| 20181106 | Pol32 | 13 | 90.77 [41.43, 140.11] |
| 20190329 | Pol3 | 21 | 131.43 [75.22, 187.65] |
| 20190403 | Pol3 | 23 | 80 [47.31, 112.69] |
| 20181116 | Histone H3 | 60 | 153.67 [114.78, 192.55] |

**Table S8** – Results showing the track durations of all experiments. Histone H3 (20180511) was used for the ML training data set.
